# Metabolic engineering of *Yarrowia lipolytica* for higher fatty alcohol production

**DOI:** 10.64898/2026.10.21.760811

**Authors:** Ebin Joseph, Fakhraddin Eminli, Payman Tohidifar, Sujit Jagtap, Christopher V. Rao

**Author notes:** These authors contributed equally to this work (Ebin Joseph, Fakhraddin Eminli). Corresponding author., Address: 600 S. Mathews Ave., Urbana, IL 61801, USA.

## Abstract

Fatty alcohols are versatile chemicals with growing demand across a broad range of industries, yet conventional production methods pose environmental challenges. Microbial biosynthesis offers a promising alternative, if hosts can be engineered to achieve high product titers. The industrially relevant oleaginous yeast Yarrowia lipolytica is well suited to this challenge because of its natural capacity for lipid metabolism. Here, we engineered Y. lipolytica W29 for fatty alcohol production by disrupting pathways involved in lipogenesis and fatty alcohol oxidation, identifying fatty alcohol oxidation as a major constraint on product accumulation. Initial engineering yielded fatty alcohol titers of 2.5 g/L from 40 g/L glucose. Subsequent optimization of media supplements and nutrient concentrations, combined with deletion of the citrate transporter CEX1 and addition of 1% Tergitol to the extractive culture medium, increased titers to 10.9 g/L in shake flasks with 80 g/L glucose and a carbon-to-nitrogen ratio of 60. Scaling the process to fed-batch bioreactor cultivation further increased titers to 19.4 g/L. These results demonstrate that combining metabolic engineering with medium and process optimization enables high-titer fatty alcohol production in *Y. lipolytica*.

**Highlights:**

- Increased fatty alcohol titers by disrupting lipogenesis, fatty alcohol oxidation, and citrate export
- Dodecane and Tergitol addition increased flask fermentation titers to 10.7 g/L
- Fed-batch fermentation in bioreactors was performed with Tergitol to 19.4 g/L

## 1. Introduction

Fatty alcohols are valuable oleochemicals with an aliphatic acyl chain of more than eight carbon atoms and a terminal hydroxyl group. The global fatty alcohol market was valued at $6.05 billion in 2024 and is projected to reach $8.63 billion by 2032 (Research and Markets, 2026). These compounds are widely used in detergents, cosmetics, plasticizers, and solvents, and are also being investigated as precursors to sustainable aviation fuels via alcohol-to-jet pathways (He et al., 2018; Nooto et al., 2025).

Industrial fatty alcohol production currently relies on either petrochemical or biomass-derived feedstocks (Ebrahimpourboura et al., 2026). In petrochemical processes, olefins are hydroformylated to aldehydes and then hydrogenated to fatty alcohols. Alternatively, fatty alcohols can be produced via the Ziegler process (Zhang et al., 2024). Biomass-derived fatty alcohols are commonly produced from vegetable oils by catalytic hydrogenation of fatty acids (Ali et al., 2019). Although these feedstocks are renewable, the chemical industry relies heavily on palm kernel and coconut oils. The large-scale cultivation of these crops raises environmental concerns, including deforestation, biodiversity loss, and, in some cases, relatively high greenhouse gas emissions (Meijaard et al., 2020; Ostfeld and Reiner, 2024; Shah et al., 2016). These limitations motivate the development of more sustainable production methods, including biological fatty alcohol synthesis.

In many organisms, fatty acyl-CoA reductases (*FAR*s) are the primary enzymes responsible for fatty alcohol biosynthesis (Domergue et al., 2010; Moore and Snyder, 1982; Teerawanichpan et al., 2010; Willis et al., 2011). *FAR*s are commonly classified into two groups based on their products. Aldehyde-forming *FAR*s reduce fatty acyl-CoA to a fatty aldehyde, which is then converted to a fatty alcohol by a second enzyme. In contrast, alcohol-forming *FAR*s catalyze the complete reduction of fatty acyl-CoA to a fatty alcohol within a single enzyme (Vioque and Kolattukudy, 1997). An alternative pathway uses carboxylic acid reductase (*CAR*) to produce fatty aldehydes directly from free fatty acids rather than from activated fatty acyl-CoA substrates (Akhtar et al., 2012).

Several studies have investigated the heterologous expression of fatty alcohol biosynthetic genes in microbial hosts to achieve industrially relevant titers and yields (Naccarato et al., 1974; Runguphan and Keasling, 2014; Zheng et al., 2012). In *Escherichia coli*, Liu and coworkers achieved a fatty alcohol titer of 6.33 g L ¹ from glycerol and glucose under fed-batch conditions by deleting *tesA*, *tesB*, and *tesC* to induce fatty acid starvation and eliminating key competing pathways (Liu et al., 2016). In the model yeast *Saccharomyces cerevisiae*, d’Espaux and coworkers combined push-pull-block engineering with the evaluation of several *FAR*s and found that the mouse *FAR* performed best. The resulting strain produced 6.0 g L ¹ of fatty alcohols in fed-batch cultivation, with a yield of 58 mg g ¹ glucose (d’Espaux et al., 2017). Other studies have focused on controlling product chain-length distribution. For example, expression of *acr1* in *E. coli* resulted in up to 75% selectivity toward C12 and C14 fatty alcohols (Zheng et al., 2012).

Oleaginous yeasts are particularly promising hosts because they can accumulate more than 20% of their cellular dry weight as triacylglycerols in intracellular lipid bodies under nutrient limiting conditions (Holdsworth and Ratledge, 1988; Lamers et al., 2016; Ratledge and Wynn, 2002). Their ability to generate and store large pools of fatty acids makes them attractive platforms for producing biodiesel and other high-value fatty acid-derived chemicals, including fatty alcohols (Louhasakul et al., 2018; Ratledge, 1994). Oleaginous yeasts investigated for fatty alcohol production include *Lipomyces starkeyi*, *Rhodotorula toruloides*, and *Yarrowia lipolytica* (Cordova et al., 2020; Fillet et al., 2015; McNeil and Stuart, 2018; Soong et al., 2021; Wang et al., 2016a). In *L. starkeyi*, multiple integrations of *mhFAR*, which encodes a highly active *FAR* from *Marinobacter*, yielded 770 mg L ¹ of fatty alcohols in flask cultivation and 4.2 g L ¹ after scale-up to fed-batch bioreactors (Wang et al., 2020). Expression of the same enzyme in *R. toruloides* produced 8 g L ¹ from sucrose after 75 h of fed-batch cultivation (Fillet et al., 2015).

As the best-characterized oleaginous yeast, *Y. lipolytica* has considerable potential for fatty alcohol production. It is generally recognized as safe (GRAS) status, tolerance to chemical and physical stressors, and an extensive genetic toolkit make it an attractive platform for industrial bioproduction (Park and Ledesma-Amaro, 2023). Lipid contents approaching 70-80% of cellular dry weight have been reported in some strains, further supporting its potential for oleochemical production (Qiao et al., 2015). In a diacylglycerol acyltransferase 1-overexpressed *Y. lipolytica* L36 strain with a mutated *Mga2* regulator carrying two integrated copies of *mhFAR*, Cordova et al. achieved fatty alcohol titers of 1.5 g L ¹ in fed-batch flask cultivation and 5.8 g L ¹ in a 2-L bioreactor (Cordova et al., 2020). Despite these advances, further improvements in pathway engineering and cultivation conditions are required to increase fatty alcohol titers and yields in this host.

In this study, we systematically deleted key genes involved in fatty alcohol and fatty acyl-CoA metabolism in a fatty-alcohol-producing *Y. lipolytica* strain and further enhanced production by optimizing a defined growth medium (**Figure 1**). Disruption of triacylglycerol synthesis and fatty alcohol oxidation generated a high-producing chassis strain. Deletion of the citrate exporter gene further supported production under nitrogen-limited conditions, yielding a fatty alcohol titer of up to 7.5 g L ¹ in minimal medium. The addition of an extractive overlay and a surfactant increased the titer to 10.87 g L ¹, with a yield of 13.6 mg g ¹ glucose, the highest flask-scale titer reported to date, to our knowledge. Upon scale-up to a bench-scale bioreactor, the engineered strain reached a peak titer of 19.35 g L ¹, representing, to our knowledge, the highest reported titer for microbial fatty alcohol production.

**Figure 1.**
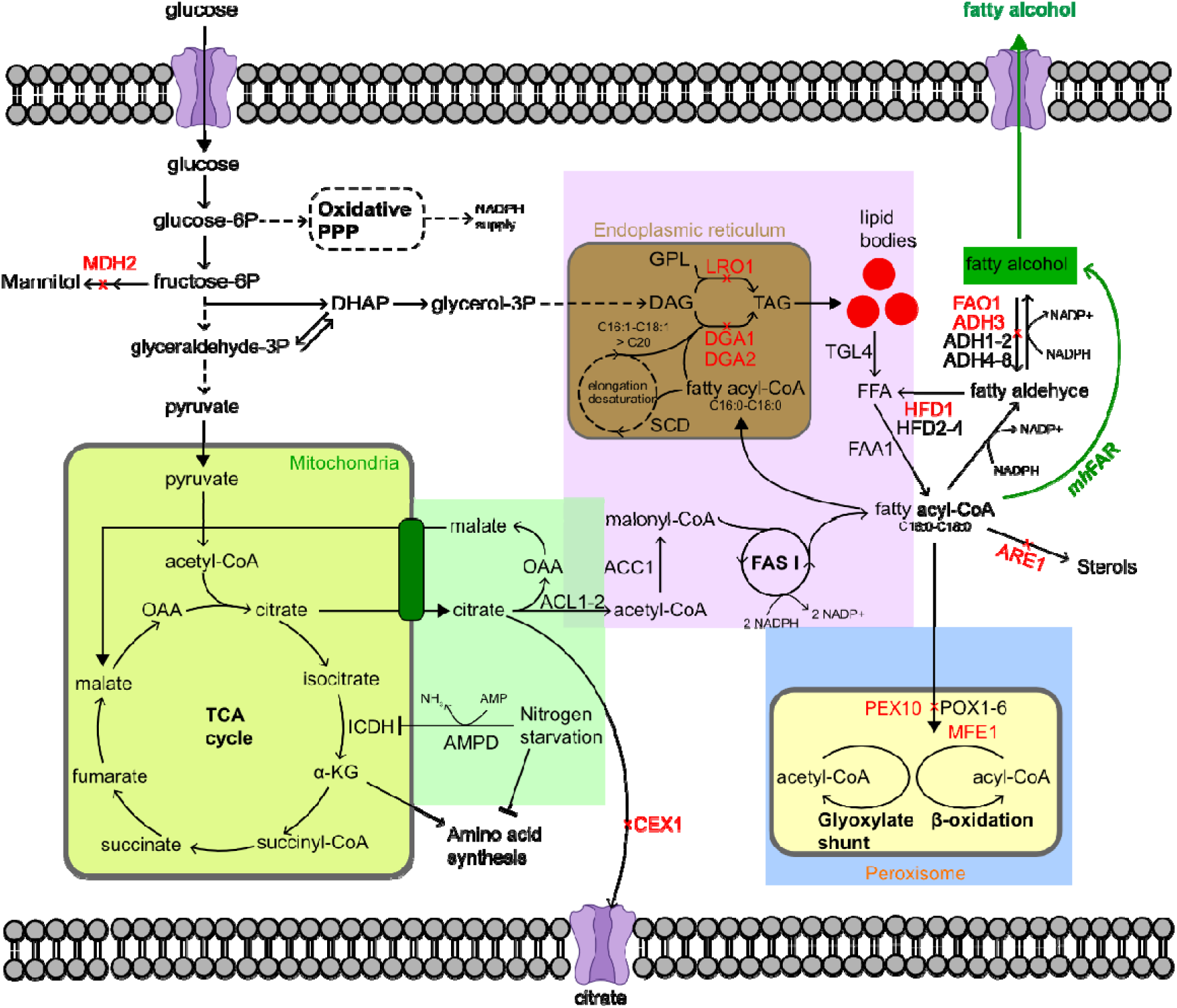
Schematic of fatty alcohol production pathway in *Y. lipolytica* and genetic targets explored in this study. Red genes denote gene deleted in this study and green denotes *mhFAR* heterologous expression. Abbreviations: *MDH2* – mannitol dehydrogenase; *LRO1* – lecithin cholesterol acyltransferase; *DGA1* – diacylglycerol acyltransferase 1; *DGA2* – diacylglycerol acyltransferase 2; *SCD* – stearoyl-CoA desaturase; *TGL4* – Triacylglycerol Lipase 4; *HFD1-4* – fatty aldehyde dehydrogenase 1-4; *FAA1* – fatty acyl-CoA synthetase; *FAO1* – fatty alcohol oxidase; *ADH1-8* – alcohol dehydrogenase 1-8; *ARE1* – acyl-CoA:sterol acyltransferase; *PEX10* – peroxisome biogenesis factor 10; *POX1-6* – acyl-coenzyme A oxidase; *MFE1* – multifunctional enzyme 1; *ACC1* – acetyl-CoA carboxylase; *ACC1* – acetyl-CoA carboxylase; *ACC1* – acetyl-CoA carboxylase; *ACL1-2* – ATP-citrate lyase; *CEX1* – citric acid exporter 1; *AMPD* – adenosine monophosphate deaminase; *ICDH* – isocitrate dehydrogenase; *mhFAR* – *M. hydrocarbonoclasticus* fatty acyl-CoA reductase.

## 2. Materials and Methods

### 2.1. Strains and culture conditions

All strains were isogenic derivatives of *Y. lipolytica* W29 (ATCC 20460) and are listed in **Table 1**. YPD medium (10 g/L yeast extract, 20 g/L peptone, and 20 g/L glucose) was used for routine growth. Unless otherwise specified, experiments were performed in YNB medium containing 1.7 g/L YNB without amino acids or ammonium sulfate, 15 g/L ammonium sulfate, 100 mM potassium phosphate (pH 6.8), and 40 g/L glucose, corresponding to a C:N ratio of 5. For experiments requiring a higher glucose concentration, glucose was increased from 40 to 80 g/L. Under low-nitrogen conditions with 80 g/L glucose, the ammonium sulfate concentration was reduced to 2.5 or 1.5 g/L to achieve C:N ratios of 60 or 100, respectively. For the bioreactor experiments, the following growth medium was employed: 1.7 g/L YNB without amino acids or ammonium sulfate, 3.75 g/L ammonium sulfate, 100 mM potassium phosphate (pH 5.6), and 120 g/L glucose, corresponding to a C:N ratio of 60.

Cells were revived by plating on YPD. Three colonies were then inoculated into separate 14 mL round-bottom tubes containing 2 mL of YPD. The cultures were grown for 24 h, washed with phosphate-buffered saline, and then again with deionized water. The supernatant was then replaced with YNB medium supplemented with the specified concentrations of ammonium sulfate and glucose. The seed cultures were then used to inoculate the 125 mL flasks (with small baffles) with 25 mL of media at a seed optical density of 0.2. The fermentation experiments were conducted on orbital shakers operating at 250 rpm and 30 °C. For extractive fermentations, 16% (v/v) dodecane was added after 24 h of growth. Tergitol was added after 24 h of growth at concentrations of 0.1% or 1% (v/v) when employed.

### 2.2. Plasmid Construction

The *mhFAR* expression cassette was constructed using the modular Golden Gate toolkit for *Y. lipolytica* (Larroude et al., 2019). The fatty-acyl CoA reductase gene (*mhFAR*) from *Marinobacter hydrocarbonoclasticus* was codon-optimized for *Y. lipolytica* (**Table S4**) and synthesized by Twist Bioscience (South San Francisco, CA, USA). The codon-optimized *mhFAR* gene was then cloned into the vector GGE0029 (Addgene #120730) under the control of the pTEF-4UAS promoter (Addgene #120751) and *LIP2* terminator (Addgene #120774). The expression cassette was cloned downstream of a hygromycin resistance cassette (Addgene #120736) with flanking LoxP sites. The complete construct was flanked by 1-kb upstream and downstream homology arms to facilitate integration at the A08 locus.

CRISPR-Cas9 knockout plasmids were constructed using JME4580 (Addgene plasmid #129659) and JME4599 (Addgene #129660) (Larroude et al., 2020). The sgRNAs were selected via CHOPCHOP for proximity to the start codon, high editing efficiency, and zero or minimal mismatches (Labun et al., 2019). Two different strategies were employed for deletions. The first strategy was a clean deletion of the genes, in which a 40 bp deletion template was assembled by overlap extension PCR and transformed into cells alongside a CRISPR plasmid containing the appropriate sgRNA. Alternatively, a dual-cut CRISPR plasmid system was also utilized. Two sgRNA sequences were chosen via CHOPCHOP (Labun et al., 2019), spanning most of the gene coding DNA sequence, one targeting a cut site close to the start codon and the other one close to the stop codon. Both sgRNAs were cloned into the base CRISPR vectors with each expressing a different known antibiotic, nourseothricin (400 μg/mL) or hygromycin (250 μg/mL), and co-transformed into yeast cells with lithium-acetate transformation method as described below.

### 2.3. Strain construction

To create the *mhFAR* expression strain, the *mhFAR* expression plasmid was first digested with NotI, gel-purified, and then transformed into cells along with the CRISPR-Cas9 plasmid containing the A08 target sgRNA, using the transformation protocol described below. The A08 site was selected as the target for integrating the *mhFAR* expression cassette, because it reportedly supports high expression and is not proximal to any known gene. After verification of successful transformants, pMCS-UAS1B_16_-TEF-Cre plasmid (Blazeck et al. 2013) was transformed into the cells to remove the antibiotic resistance cassette.

The lithium acetate method, with minimal modifications, was used for yeast transformations. Briefly, a transformation master mix was prepared with the following components: 200 mM lithium acetate, 100 mM dithiothreitol, 1x TE buffer (10 mM Tris and 1 mM EDTA, pH 8), 2 mg/mL ssDNA, and 40% PEG4000. The cells were suspended in the master mix, and the desired plasmids or purified deletion templates were added in quantities of at least 2 µg in a PCR tube. The cells were well mixed by pipetting and heat-shocked at 39°C for 30 min in either a water bath or a thermocycler. The tubes were then cooled to room temperature and centrifuged to pellet the cells. The master mix was then removed and replaced with YPD liquid media. The cells were either recovered for 2-4 hours at 30°C or plated directly on antibiotic-containing plates (YPD agar). The plates were incubated at 30°C for 3-4 days until colonies appeared. The transformants were cultured in non-selective medium to cure the transformed plasmids.

### 2.4. Analytical Measurements

Cell growth was quantified by measuring optical density at 600nm using a Biocrom WPA Biowave CO8000 Cell Density Meter (Cambridge, UK). The cuvette was blanked with water, and cells were added, yielding a dilution factor of 50 or 100 to stay within the linear range of the transmittance readings.

Fatty alcohol concentrations were measured using a gas chromatograph equipped with a flame ionization detector. 500 µL of culture was aliquoted into 2 mL microcentrifuge tubes. An equal volume of ethyl acetate was added to tubes designated for product extraction, along with approximately 200 µL of acid-washed glass beads. The tubes were secured on a vortex mixer and run at maximum speed for 1 h, followed by centrifugation at maximum speed for 10 minutes. 1 µL of 1-nonadecanol was added to a glass insert at a final concentration of 100 mg/L, and 99 µL of the top organic phase was added to a GC vial containing a glass insert and mixed. The organic overlay samples were diluted 20-fold in ethyl acetate. Prior to analysis, the vials were stored at –20 °C. An Agilent 7890A gas chromatography-flame ionization detector system equipped with a DB-FATWAX column (30 m x 320 μm x 0.25 μm) (Santa Clara, CA) was used for fatty alcohol quantification. Airflow to the detector, held at 280 °C, was supplied at 450 mL/min, alongside a 40 mL/min H_2_ fuel flow rate. The oven was held at 50 °C for 1 minute, then ramped to 150 °C at 25 °C/min. The ramp rate was reduced to 4 °C/min to heat the oven to 250 °C, and the oven was held at that temperature for 5 minutes.

Sugar, sugar alcohol, and organic acids were measured by high-performance liquid chromatography (HPLC). 1.5 mL of culture was centrifuged at maximum speed for 10-15 minutes, and 20 μL of the supernatant was added to HPLC tubes and mixed with 180 μL of autoclaved water. Concentrations were determined using a Shimadzu high-performance liquid chromatography system equipped with a RID-10A refractive index detector (Kyoto, Japan) and a Bio-Rad Aminex HPX-87H column (Hercules, CA). 5 mM H_2_SO_4_ served as the mobile phase. Because glucose co-migrated with the phosphate buffer in the media, the analysis was performed at 30 °C and a flow rate of 0.3 mL/min. These run parameters allowed for quantification of mannitol and arabitol as well. To quantify citrate production, a column temperature of 65 °C and a flow rate of 0.6 mL/min were used. Authentic standards were used to generate calibration curves for these compounds.

### 2.5. Bioreactor operation

The bioreactor experiments were conducted using the Eppendorf DASbox Mini Bioreactor System (Hamburg, Germany). pH probes were calibrated with stock solutions of pH 4 and 7. pH and DO probes were attached to the headplate, along with the sample port, base addition port, septum, and one marine pitch-blade impeller. The bioreactors were filled with 75 mL of production media. DO calibration was performed overnight at a sparging rate of 26 sLph and 800 rpm. 3 drops of pre-emptive antifoam 204 (Sigma Aldrich A6426) were added before inoculation. Samples were inoculated to an initial OD of 0.2. A dodecane overlay at 16% and Tergitol at 0.1% were added 24 h after inoculation. DO was set to 30%, and pH was set to 5.6 with a 0.5 pH deadband by adding 33% ammonium hydroxide. Temperature was set to 30°C. If excessive foaming was observed, antifoam was added as needed. 2.5 mL samples were collected daily for analysis. For fed-batch operation, 80 g/L glucose was added in 10 mL increments. As needed, culture volume was adjusted to keep the total fermentation volume constant.

## 3. Results

### 3.1. Establishing fatty alcohol production in *Y. lipolytica* W29

To first produce fatty alcohols, we expressed a heterologous fatty acyl-CoA reductase (*mhFAR*) from *Marinobacter hydrocarbonoclasticus*. Although several fatty acyl-CoA reductases from diverse species have been explored in the literature, we chose *mhFAR* because it has previously been shown to produce fatty alcohols in yeast at relatively high titers (Cordova et al., 2020). We then tested the resulting strain in shake flasks in YNB medium containing 40 g/L glucose with a C:N ratio of 5 (**Figure 2**). This strain reached a peak fatty alcohol titer of 63 mg/L after 72 h of growth. The fatty alcohols consisted of a mixture of hexadecanol, octadecanol, and oleyl alcohol, with octadecanol as the main product (70-80%), followed by hexadecanol (20-30%) (**Figure S1**). In addition to fatty alcohols, we observed the production of citrate, mannitol, and arabitol, though production was similar in both wild-type (W29) and *mhFAR*-expressing strains (**Figure 2**). In addition, *mhFAR* expression reduced growth by approximately 40% relative to the wild type. This suggests that fatty alcohols impose a metabolic burden on cells or are toxic to cell growth.

**Figure 2.**
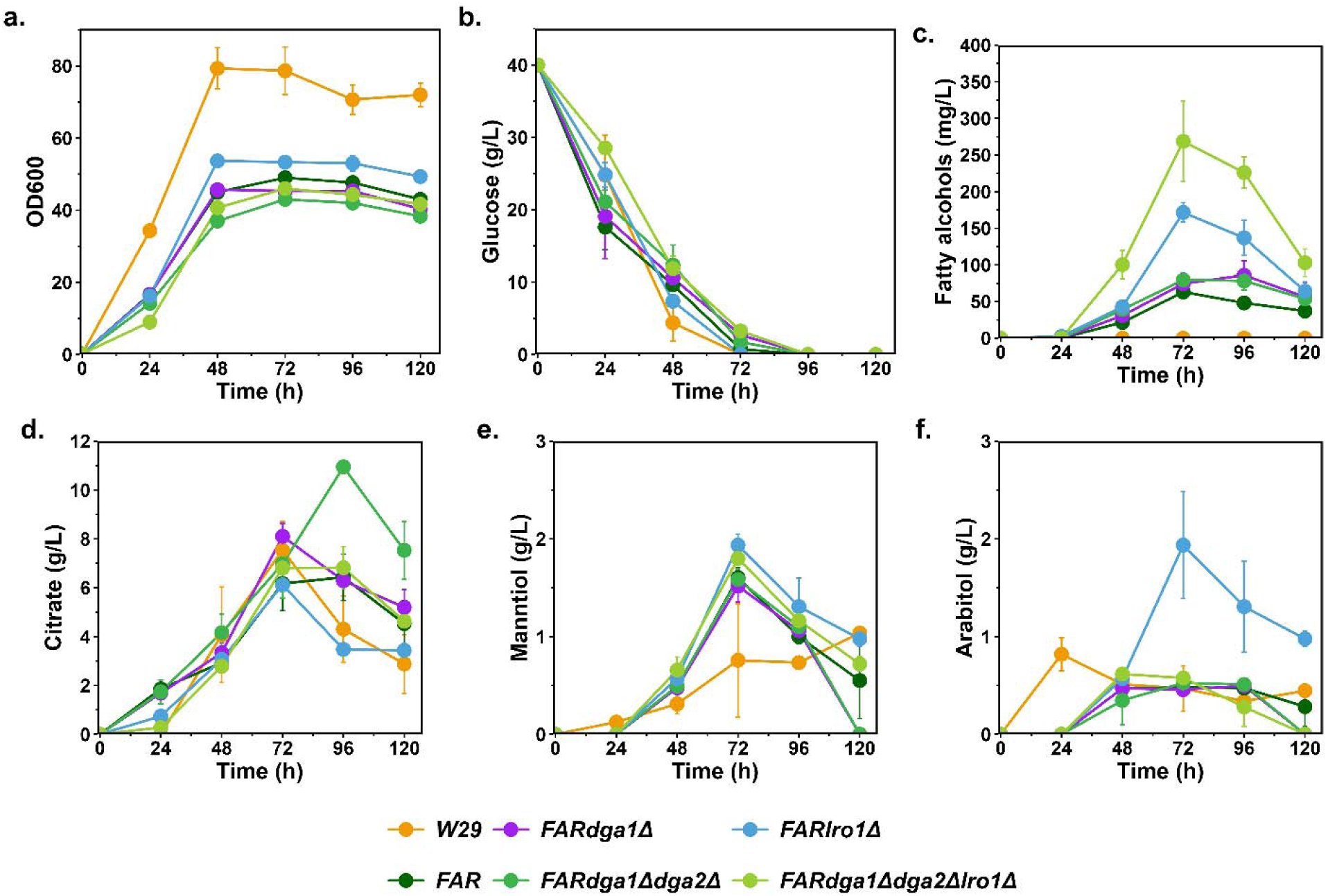
Growth and fermentation profiles of triacylglycerol-deficient strains. Growth experiments were performed in shake flasks containing YNB medium with 40 g/L glucose (C/N 5) using shake flasks. Legend: (a.) cell growth (OD600), (b.) glucose consumption, (c.) total fatty alcohol production, (d.) citrate production, (e.) mannitol production, and (f.) arabitol production. Error bars denote the standard deviation from three biological replicates.

### 3.2. Eliminating competition for potential fatty alcohol precursors increases fatty alcohol titers

We next sought to increase the availability of fatty acyl-CoA substrates by eliminating triacylglycerol synthesis. In *Y. lipolytica*, three enzymes are known to support TAG synthesis: diacylglycerol acyltransferase 1 (*DGA1*), diacylglycerol acyltransferase 2 (*DGA2*), and lecithin cholesterol acyltransferase (*LRO1*). We tested whether deleting these genes would increase fatty alcohol titers (**Figure 2**). We first deleted *DGA1* and found that peak fatty alcohol titers increased to 86 mg/L when cells were cultured in YNB medium containing 40 g/L glucose at a C:N ratio of 5. We then deleted *DGA2* in this strain (*dga1*Δ*dga2*Δ) but observed no change in peak fatty alcohol titers in the same growth medium: 79 mg/L. When we also deleted *LRO1* (*dga1*Δ*dga2*Δ*lro1*Δ), peak fatty alcohol titers nearly quadrupled to 269 mg/L in the same growth medium. These results suggest that *LRO1* competes with *mhFAR* for fatty acyl-CoA substrates. In support of this hypothesis, fatty alcohol titers were 171 mg/L in a strain deleted for *LRO1* alone. We also tested the effect of deleting acyl-CoA:sterol acyltransferase (*ARE1*) in the *dga1*Δ*dga2*Δ*lro1*Δ strain but observed a small reduction in fatty alcohol production (**Figure S2**). Therefore, we did not explore *ARE1* further.

### 3.3. Eliminating fatty alcohol catabolism increases fatty alcohol titers

*Y. lipolytica* can metabolize fatty alcohols, which are first oxidized to fatty aldehydes and then to fatty acids, which can subsequently be metabolized via beta-oxidation. This reaction is catalyzed by the irreversible fatty alcohol oxidase (*FAO1*). Deletion of *FAO1* in the *mhFAR*-expressing strain significantly increased fatty alcohol titers (**Figure 3**), with a peak titer of 2.1 g/L. This represents a 32-fold increase over the base *mhFAR*-expressing strain. These results indicate that fatty alcohols are produced but are quickly catabolized in the base strain. However, even in the *fao1*Δ mutant, fatty alcohols are catabolized once glucose is depleted from the growth medium. Knocking out *FAO1* in the triacylglycerol-deficient strain (*dga1*Δ*dga2*Δ*lro1*Δ) yielded peak fatty alcohol titers of 2.3 g/L, a 10% increase compared to the *fao1*Δ mutant.

**Figure 3.**
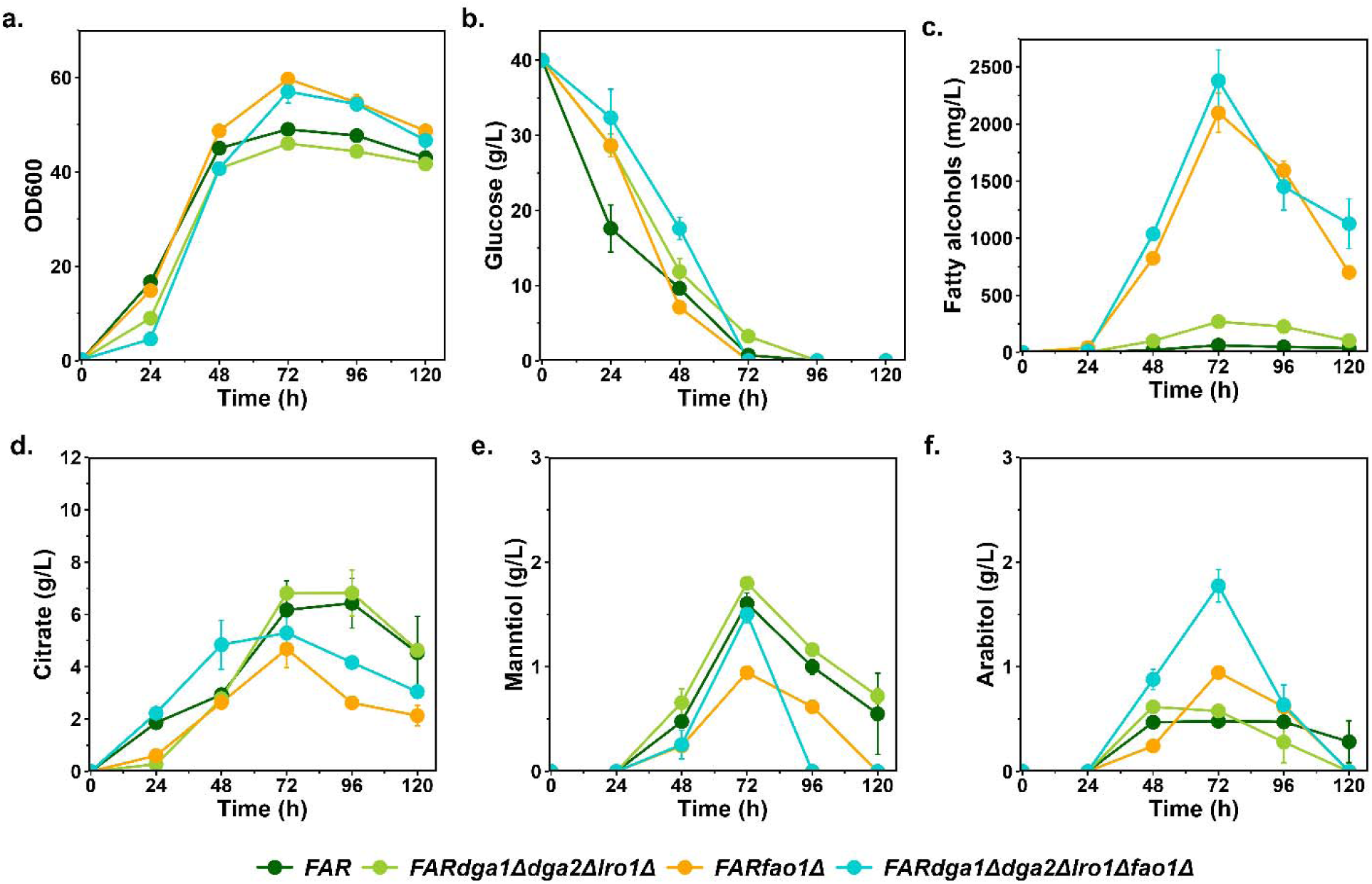
Effect on *FAO1* knockout fatty alcohol production. Growth experiments were performed in shake flasks containing YNB medium with 40 g/L glucose (C/N 5) using shake flasks. Legend: (a.) cell growth (OD600), (b.) glucose consumption, (c.) total fatty alcohol production, (d.) citrate production, (e.) mannitol production, and (f.) arabitol production. Error bars denote the standard deviation from three biological replicates.

Given *FAO1*’s prominent role in fatty alcohol production, we focused on other genes that oxidize fatty alcohols to fatty aldehydes. We focused on alcohol dehydrogenase 3 (*ADH3*), which oxidizes fatty alcohols. However, deleting *ADH3* in the *FAO1* strain resulted in no fatty alcohol production (data not shown). Therefore, we did not further explore the deletion of other alcohol– or aldehyde-dehydrogenase genes in this study. We also tested whether disrupting beta-oxidation would improve fatty alcohol production. We tested deleting *PEX10* and *MFE1*, as both have previously been shown to improve fatty alcohol production in *Y. lipolytica* (Dahlin et al., 2019; Rutter and Rao, 2016; Somasundaram et al., 2026). *PEX10* is responsible for peroxisome biogenesis, whereas *MFE1* is a bi-functional enzyme that catalyzes the second and third steps of β-oxidation (Haddouche et al., 2011; Wei et al., 2021). Both genes were deleted in the *dga1*Δ*dga2*Δ*lro1*Δ*fao1*Δ strain. Deleting *PEX10* hindered growth and reduced peak fatty alcohol titers by 63% compared to the *dga1*Δ*dga2*Δ*lro1*Δ*fao1*Δstrain (**Figure S3**). The *MFE1* deletion was also deleterious, resulting in a 23% loss of titers (**Figure S4**). Because neither gene deletion improved titers, they were not pursued further.

### 3.4. Optimizing medium composition and eliminating secreted byproducts improves fatty alcohol production

We next investigated different medium compositions to determine whether they could increase fatty alcohol titers. As a first step, we increased the glucose concentration from 40 g/L to 80 g/L in YNB medium. This doubled the fatty alcohol titers to 4.8 g/L in our best strain, *dga1*Δ*dga2*Δ*lro1*Δ*fao1*Δ (**Figure S5**). However, when we increased the glucose concentration to 120 g/L, the titers did not increase further (**Figure S6**). We also attempted to supplement the growth medium with yeast extract or alter the media formulation. While these changes to the growth medium improved the growth rate, the fatty alcohol titers did not increase substantially and, in many cases, decreased compared with results obtained in YNB medium.

Many studies on fatty alcohol production in yeast use an organic overlay, often dodecane, to sequester fatty alcohols in the organic phase (Cordova et al., 2020; Wang et al., 2020). This approach is thought to limit fatty alcohol toxicity and improve their solubility. Our previous results did not include an organic overlay. Therefore, we tested whether adding dodecane would improve fatty alcohol titers. Peak titers were unchanged (**Figure S7**). However, adding dodecane prevented fatty alcohol catabolism. When we determined the location of fatty alcohols, we found that they were almost entirely intracellular in the absence of dodecane but were primarily extracellular in its presence. The elimination of fatty alcohol catabolism likely results from sequestering fatty alcohols outside the cell, so intracellular degradation cannot occur. Further results involving dodecane are discussed in the next section.

We next tested the effect of altering the C:N ratio of the growth medium by varying the amount of ammonium sulfate added to YNB medium containing 80 g/L glucose (**Figure 4**). Three different C:N ratios were tested: 5, 60, and 100. The best production was obtained at a C:N ratio of 60, with peak fatty alcohol titers of 5.5 g/L. This was slightly better than the results obtained at a C:N ratio of 5, where the peak fatty alcohol titers were 5.0 g/L. However, peak titers were significantly reduced to 2.4 g/L at a C:N ratio of 100.

**Figure 4.**
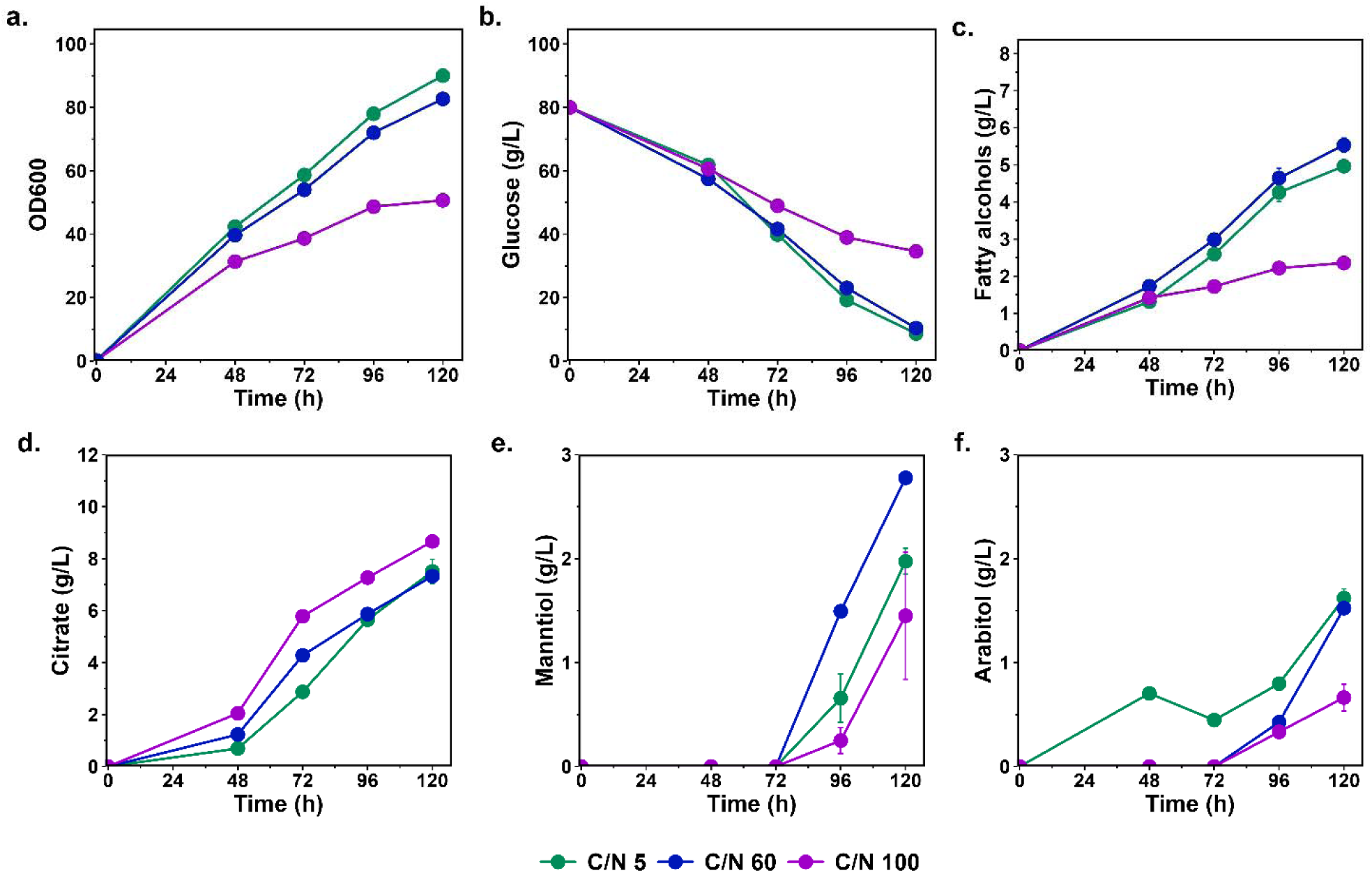
Effect of C/N ratio on fatty alcohol production. Growth experiments were performed using the *mhFAR*-expressing *dga1*Δ*dga2*Δ*lro1*Δ*fao1*Δ strain in shake flasks containing YNB medium with 80 g/L glucose at the specified C/N ratio. Legend: (a.) cell growth (OD600), (b.) glucose consumption, (c.) total fatty alcohol production, (d.) citrate production, (e.) mannitol production, and (f.) arabitol production. Error bars denote the standard deviation from three biological replicates.

Under all tested conditions, we observed significant citrate secretion that increased with the C:N ratio. Because citric acid is a source of cytoplasmic acetyl-CoA via ATP citrate lyase, we sought to eliminate citrate secretion by deleting the citrate exporter, *CEX1*. When we deleted *CEX1* in the *dga1*Δ*dga2*Δ*lro1*Δ*fao1*Δ strain (**Figure 5**), we eliminated citrate secretion as expected. In addition, peak fatty alcohol titers increased by 21% and 94% at C:N ratios of 60 and 100, respectively. However, they increased by only 1% at a C:N ratio of 5. The best performance was achieved at a C:N ratio of 60, with a peak titer of 6.7 g/L. Although the largest increases were observed at a C:N ratio of 100, the peak titers were 4.6 g/L, far less than those obtained at a C:N ratio of 60.

**Figure 5.**
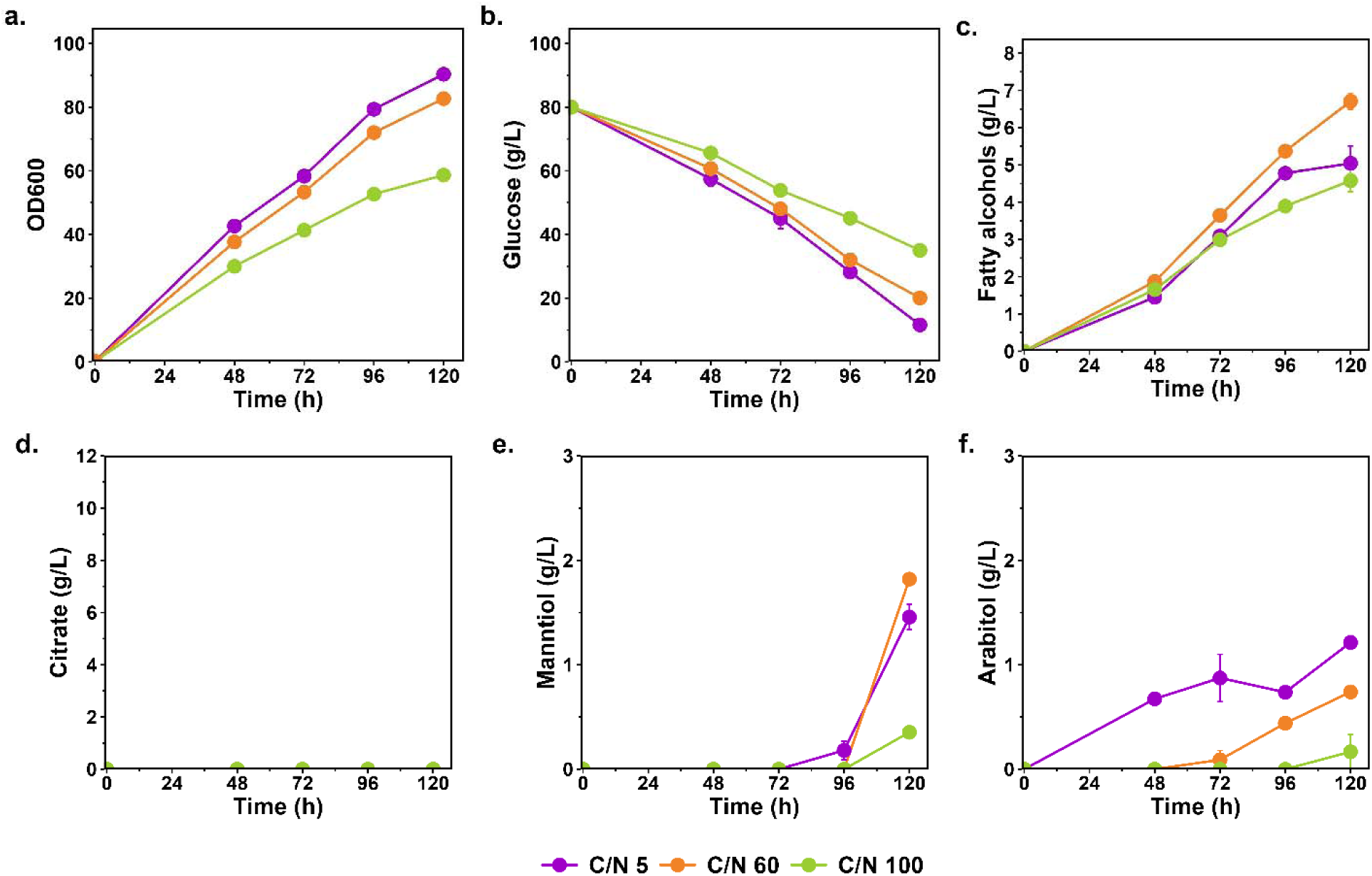
Effect of C/N ratio test for fatty alcohol production in strain deficient in citrate export. Growth experiments were performed using the *mhFAR*-expressing *dga1*Δ*dga2*Δ*lro1*Δ*fao1*Δ*cex1*Δ strain in shake flasks containing YNB medium with 80 g/L glucose at the specified C/N ratio. Legend: (a.) cell growth (OD600), (b.) glucose consumption, (c.) total fatty alcohol production, (d.) citrate production, (e.) mannitol production, and (f.) arabitol production. Error bars denote the standard deviation from three biological replicates.

We also attempted to eliminate polyol secretion by deleting the gene encoding mannitol dehydrogenase (*MDH2*) in the *dga1*Δ*dga2*Δ*lro1*Δ*fao1*Δ strain (**Figure S8**). Deleting *MDH2* eliminated both mannitol and arabitol production. However, it reduced peak fatty alcohol titer. We also tested a strain where both citrate and polyol secretion were eliminated (*dga1*Δ*dga2*Δ*lro1*Δ*fao1*Δ*cex1*Δ*mdh2*Δ). While both citrate and polyol secretion were eliminated in this strain, peak fatty alcohol titers were less than those obtained with the *dga1*Δ*dga2*Δ*lro1*Δ*fao1*Δ and *dga1*Δ*dga2*Δ*lro1*Δ*fao1*Δ*cex1*Δ strains (**Figure S8**). Since the best performance was obtained with the *dga1*Δ*dga2*Δ*lro1*Δ*fao1*Δ*cex1*Δstrain, we used this for the remainder of our investigations.

Previous results have demonstrated that the addition of tergitol NP-40 increases fatty alcohol titers in the oleaginous yeast *Rhodotorula toruloides* (Liu et al., 2020). Therefore, we tested whether the addition of tergitol would also increase fatty alcohol production in the *dga1*Δ*dga2*Δ*lro1*Δ*fao1*Δ*cex1*Δ strain (**Figure S9**). The use of tergitol alone did not yield any significant differences in fatty alcohol titers. However, when used in conjunction dodecane, fatty alcohol titer increased from 7.2 g/L to 9.1 g/L when 0.1% (v/v) tergitol was used and 10.9 g/L when 1% tergitol (v/v) was used (**Figure 6**). Interestingly, tergitol and docadane also improved growth.

**Figure 6.**
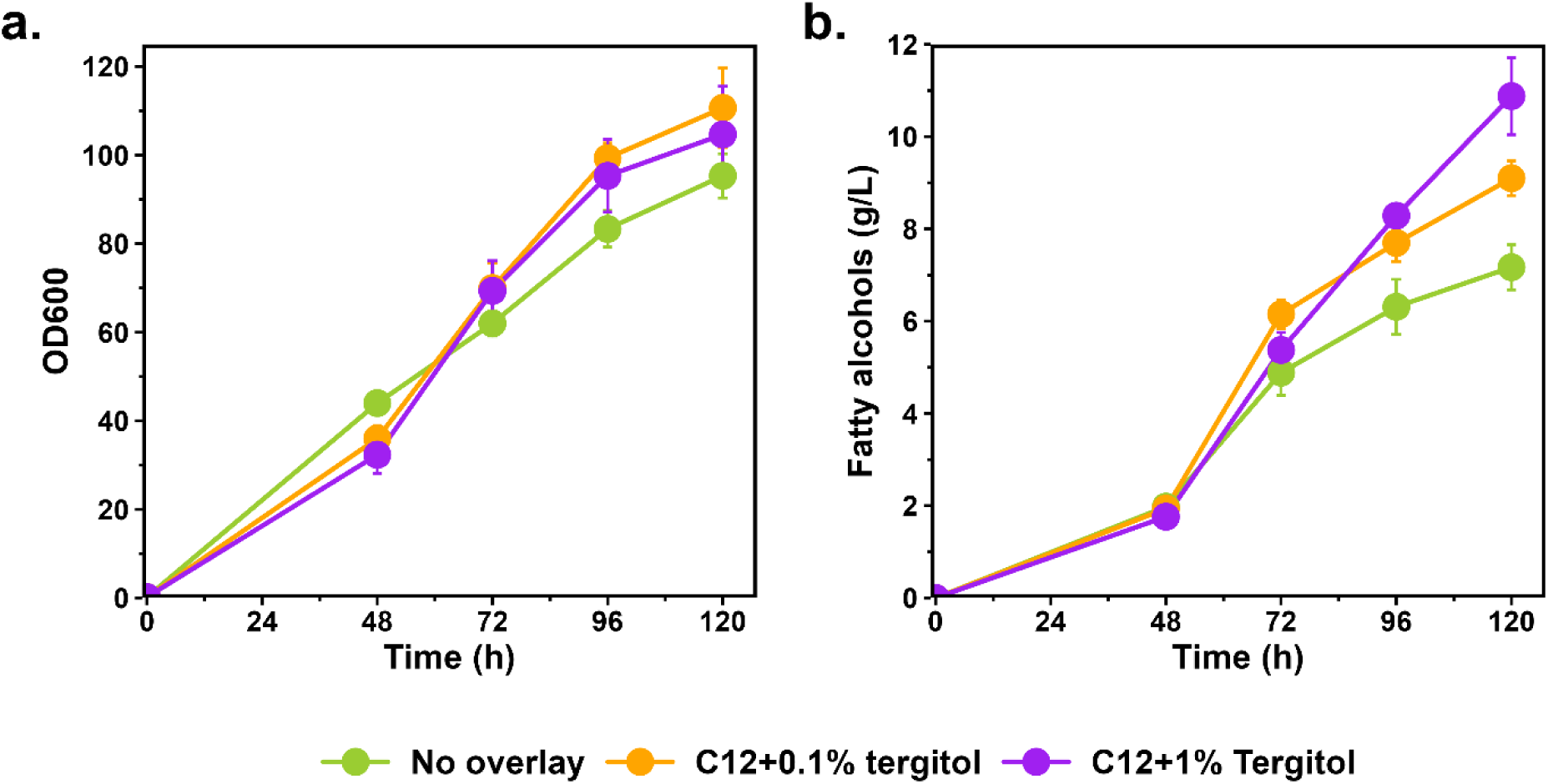
Effect of dodecane and tergitol on fatty alcohol production. Growth experiments were performed using the *mhFAR*-expressing *dga1*Δ*dga2*Δ*lro1*Δ*fao1*Δ*cex1*Δ strain in shake flasks containing YNB medium with 80 g/L glucose and a C/N ratio of 60. Legend: (a.) cell growth (OD600) and total fatty alcohol production. Error bars denote the standard deviation from three biological replicates.

### 3.5. Performance in fermenters

Our best results were obtained using the *dga1*Δ*dga2*Δ*lro1*Δ*fao1*Δ*cex1*Δstrain in YNB medium containing 80 g/L glucose, a C:N ratio of 60, 16% (v/v) dodecane, and 0.1% (v/v) tergitol. The lower tergitol concentration was used to reduce foaming. These experiments were performed in 125 mL shake flasks with a working volume of 25 mL. To further test performance, we then conducted fed-batch experiments in mini (250 mL) bioreactors with an initial glucose concentration of 120 g/L. An additional 80 g/L of glucose was added after 72 and 120 h of growth. In addition, 2.5 g/L of ammonium sulfate was added after 120 h to promote further growth. To prevent excessive foaming, we used pitch-blade impellers after initially trying Rushton impellers. Under these conditions, we achieved a peak fatty alcohol titer of 19.4 g/L after 192 h of growth (**Figure 7**). Glucose was not fully consumed, and fatty alcohol production ceased after 192 h.

**Figure 7.**
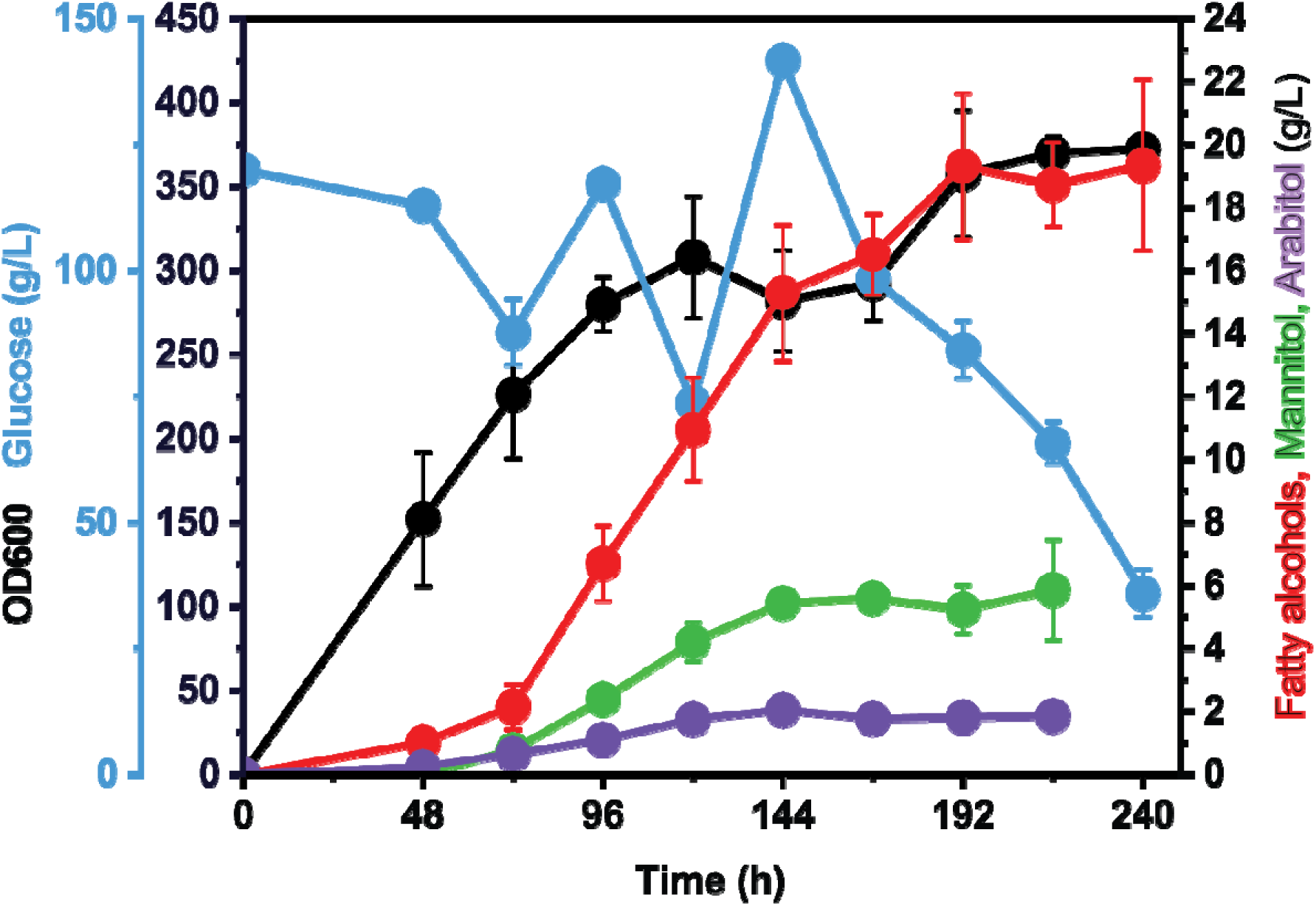
Fatty alcohol production in fermenters. Growth experiments were performed using th *mhFAR*-expressing *dga1*Δ*dga2*Δ*lro1*Δ*fao1*Δ*cex1*Δ strain in bioreactors with starting 120g/L of glucose and C/N ratio of 60. An additional 80 g/L of glucose was added at 72 h and 120 h with an additional 2.5g/L of ammonium sulfate added at 120 h. Error bars denote the standard deviation from two technical replicates.

## Discussion

*Yarrowia lipolytica* is a well-established host for producing fatty acyl-CoA-derived compounds, including fatty alcohols generated by fatty acyl-CoA reductase (FAR) activity (Sun et al., 2021; Xu et al., 2016; Zhang et al., 2019a). In this study, we combined rational metabolic engineering, medium optimization, and extractive fermentation to redirect fatty acyl-CoA toward fatty alcohol synthesis, limit product degradation, and overcome constraints associated with high intracellular product concentrations.

Our initial engineering strategy focused on reducing competition from triacylglycerol synthesis. Previous studies showed that FAR expression decreases cellular lipid content, suggesting that fatty alcohol production competes with lipid accumulation for fatty acyl-CoA (Wang et al., 2016b). Accordingly, *DGA1*, *DGA2*, and *LRO1* were disrupted to prevent TAG formation. Deletion of *DGA1* and *LRO1* increased fatty alcohol production by approximately 52%, whereas the subsequent deletion of *DGA2* provided no additional benefit. The improvement associated with *LRO1* deletion was accompanied by a modest increase in growth, although the final cell density remained within the range observed for other *mhFAR*-expressing strains. Nevertheless, combined disruption of *DGA1*, *DGA2*, and *LRO1* redirected fatty acyl-CoA away from TAG storage and produced an approximately fivefold increase in maximum fatty alcohol titer.

Preventing fatty alcohol degradation had an even greater effect on production. As a hydrocarbon-assimilating yeast, *Y. lipolytica* possesses a robust oxidative pathway involving alcohol dehydrogenases and a fatty alcohol oxidase (Gatter et al., 2014; Iwama et al., 2015). Deletion of *FAO1* produced the largest improvement among the individual genetic modifications evaluated. In the parental *mhFAR*-expressing strain, *FAO1* deletion increased the fatty alcohol titer by more than 30-fold. In the TAG-deficient background, it produced an approximately ninefold improvement under defined conditions containing 40 g L ¹ glucose. These results identify *FAO1* as a major contributor to fatty alcohol catabolism and demonstrate that blocking product oxidation is essential for achieving high titers.

The strong native capacity for fatty alcohol degradation may represent a cellular response to product toxicity. Because long-chain fatty alcohols resemble components of the lipid bilayer, their accumulation may disrupt membrane organization and increase permeability (Hu et al., 2018; Kranenburg et al., 2004; Wilbanks and Trinh, 2017). Product degradation may therefore help the cells maintain membrane homeostasis. Consistent with this possibility, fatty alcohol catabolism persisted even after *FAO1* deletion, particularly after glucose depletion. Under carbon-limited conditions, the cells appeared to consume previously accumulated fatty alcohols to support survival during stationary phase. Maintaining a preferred carbon source throughout cultivation is consequently important for both continued production and product retention.

Attempts to block additional steps in fatty alcohol oxidation revealed complex, strain-dependent effects. Deletion of *ADH3* unexpectedly abolished fatty alcohol production, possibly due to an uncharacterized role in redox balance, reversible alcohol metabolism, or pathway regulation. Deletion of HFD1 caused slight background-dependent effects on production. Therefore, further disruption of alcohol dehydrogenases or fatty aldehyde dehydrogenases was not pursued (data not shown).

We next investigated whether reducing the formation of common *Y. lipolytica* by-products could improve carbon allocation. Deletion of the citrate exporter gene *CEX1* modestly increased fatty alcohol titers, consistent with the redirection of intracellular citrate away from overflow metabolism and toward acetyl-CoA-dependent biosynthesis (Zhang et al., 2019b). In contrast, deletion of *MDH2* did not improve growth or production and eliminated both mannitol and arabitol formation. In the TAG– and *FAO1*-deficient background, *MDH2* deletion reduced growth and fatty alcohol production, and subsequent deletion of *CEX1* only partially recovered these phenotypes. Although citrate secretion represents a meaningful carbon loss, the comparatively small polyol pool may contribute to osmotic or redox homeostasis (Jagtap et al., 2021). Therefore, eliminating polyol synthesis was detrimental rather than beneficial.

Increasing substrate availability improved production only within a limited range. Raising the initial glucose concentration from 40 to 80 g L ¹ nearly doubled biomass formation and fatty alcohol production in several of the best-performing strains. However, increasing the concentration to 120 g L ¹ yielded diminishing returns, indicating that production had approached a physiological or metabolic limit. These effects may reflect limitations in intracellular storage capacity, product toxicity, declining culture viability, or the instability and metabolic burden associated with constitutive *<u>mhFAR</u>* expression.

Extractive fermentation proved an effective way to overcome these constraints. Dodecane was therefore used as an organic overlay to partition fatty alcohols away from the cells. Because *Y. lipolytica* can assimilate alkanes, exposure to dodecane may induce competing metabolic programs and increase population heterogeneity (Miranda et al., 2024). Nevertheless, adding Tergitol substantially improved product recovery. This nonionic surfactant increases membrane permeability and reduces the size of dodecane droplets, thereby increasing the interfacial area available for mass transfer. Consequently, the combination of dodecane and Tergitol produced a much greater increase in titer than cultivation without an overlay.

The strongest flask-scale production resulted achieved by combining the most beneficial genetic and process variables: the *dga1*Δ*dga2*Δ*lro1*Δ*fao1*Δ*cex1*Δ strain, an optimized carbon-to-nitrogen ratio, sufficient glucose, and extractive fermentation. Together, these interventions addressed successive production constraints by increasing precursor availability, limiting TAG formation and citrate secretion, preventing fatty alcohol degradation, and removing the product from the cellular environment. Under these conditions, the strain produced 10.87 g L ¹ fatty alcohols in shake flasks.

Scaling up to bench-scale bioreactors provided greater control over pH, dissolved oxygen, substrate feeding, and nitrogen availability. Flask cultures frequently reached pH values below 3, which may inhibit substrate utilization even when citrate export is disrupted. Maintaining the bioreactor at pH 5.6 with ammonium hydroxide alleviated acidification and provided an additional nitrogen source. Dissolved oxygen control was also critical because of the high oxygen demand of dense, aerobic cultures. The use of dodecane and Tergitol initially caused foaming and backpressure, but these problems were mitigated by increasing the vessel headspace, replacing the Rushton impeller with a marine impeller, and reducing the Tergitol concentration to 0.1%. Timely feeding of both glucose and ammonium sulfate further sustained biomass and production during extended cultivation. Although additional nitrogen altered the initial carbon-to-nitrogen ratio, it proved beneficial during the longer fed-batch process.

## 5. Conclusions

By integrating these strain– and process-level strategies, we achieved a peak fatty alcohol titer of 19.35 g L ¹ in a bench-scale bioreactor. To our knowledge, this represents the highest reported microbial fatty alcohol titer to date. More broadly, the results demonstrate that high-level fatty alcohol production requires more than increasing precursor flux. Product degradation, membrane toxicity, metabolic homeostasis, nutrient availability, oxygen transfer, and extracellular product partitioning must all be addressed. The engineered *Y. lipolytica* platform developed here provides a strong foundation for industrial fatty alcohol production and for the biosynthesis of other oleochemicals derived from fatty acyl-CoA. Future improvements should focus on stabilizing *mhFAR* expression, identifying the remaining routes of fatty alcohol catabolism, clarifying the regulatory roles of alcohol and aldehyde dehydrogenases, and developing scalable product-recovery systems that minimize surfactant use and hydrocarbon assimilation.

## 6. Author contributions: CRediT

Ebin Joseph: Conceptualization, Investigation, Methodology, Project administration, Writing – original draft, review & editing. Fakhraddin Eminli: Conceptualization, Investigation, Methodology, Project administration, Writing – original draft, review & editing. Payman Tohidifar: Conceptualization, Investigation, Methodology. Sujit Jagtap: Conceptualization, Methodology, Project administration, Writing – original draft, review & editing. Christopher Rao: Funding acquisition, Investigation, Methodology, Project administration, Resources, Writing – original draft, review & editing.

## 7. Acknowledgments

This work was funded by the DOE Center for Advanced Bioenergy and Bioproducts Innovation (U.S. Department of Energy, Office of Science, Biological and Environmental Research Program under Award Number DE-SC0018420). Any opinions, findings, and conclusions or recommendations expressed in this publication are those of the author(s) and do not necessarily reflect the views of the U.S. Department of Energy.

## 8. Data Availability Statement

Data will be made available upon request.

## 9. Declaration of Generative AI and AI-assisted technologies in the writing process

The authors used ChatGPT (OpenAI) and Grammarly as language-assistance tools to improve the clarity and readability of portions of the manuscript and to review grammar and spelling. After using these tools, the authors reviewed and edited the content as needed and take full responsibility for the content of the publication.

