## Supplementary Material for "Metabolic engineering of *Yarrowia lipolytica* for higher fatty alcohol production"

**Table S1.** List of primers used in this study.

| Primer | Sequence | Description |
| --- | --- | --- |
| FHp1 | GGGGGTCTCTCATTGGATCCTTCGGGTGTGAGTTG | For cloning P1 fragment overhangs onto 4UAS-TEFp promoter fragment |
| FHp2 | GGGGGTCTCTACGGCGATACGCGTATCGATACGC |  |
| FHp3 | GGTCTCTACGGCGATACGC | For replacing ATG start codon in P1 fragment 4UAS-TEFp right overhang with ATA basepairs |
| FHp4 | GGTCTCTTATTGGATCCTTCGGGTGTGAGTTG |  |
| FHp5 | GGTCTCTAATGATGGCCATCCAGCAGGTGC | For cloning G1 fragment overhangs onto mhFAR gene fragment |
| FHp6 | GGTCTCTTAGATTAAGCGGCCTTCTTTCGCTG |  |
| FHp7 | GGTCTCTAATAATGGCCATCC | For replacing ATG start codon in G1 fragment mhFAR left overhang with ATA basepairs |
| FHp8 | GGTCTCTTAGATTAAGCGGCC |  |
| FHp9 | GGTCTCTTCTAGTGTCTGTGGTATCTAAGC | For cloning T1 fragment overhangs onto LIP2 terminator fragment |
| FHp10 | GGTCTCTAAGCTGTCTTAGAGGAACGCATATAC |  |
| FHp11 | GGTCTCTGCCTGCGGCCGCTTCCACAGGCCTTCTGTTATATC | For amplification of A08 LHB on YL W29 strain with appropriate BsaI overhangs to construct golden gate homology arm fragment |
| FHp12 | GGTCTCTACCTTTAGTAGTCGGTGATGGTGTC |  |
| FHp13 | GGTCTCTGCTTAGAAGGGCATCAACGTGCTC | For amplification of A08 RHB on YL W29 strain with appropriate BsaI overhangs to construct golden gate homology arm fragment |
| FHp14 | GGTCTCTCGCAGCGGCCGCCTCAACCGCAAGACCCGCTG |  |
| FHp15 | CTCGAGAGCCAGCCTAAGC | Site-directed mutagenesis primers for removing BsaI site on A08 LHB fragment |
| FHp16 | AGCAAAGAAGCCGTCTGAATC |  |
| FHp17 | TTCGATTCCGGGTCGGCGCAATCACCGACTACTAACAAGAGTTTTA | For cloning A08 crRNA to BsmBI digested NATex_CrisprCas9-yl_RFP |
| FHp18 | GCTCTAAAACTCTTGTTAGTAGTCGGTGATTGCGCCGACCCGGAAT |  |
| FHp19 | ACTCAATGGCATCTACGAACG | For colony PCR of inserts into A08 locus on W29 |
| FHp20 | CGTCCAATATCCAAATGAGCG |  |
| FHp21 | TTCGATTCCGGGTCGGCGCATGGTGGAGATGATTGCTCGGGTTTTA | For cloning crRNA-DGA1_W29 to NATex_CrisprCas9-yl |
| FHp22 | GCTCTAAAACCCGAGCAATCATCTCCACCATGCGCCGACCCGGAAT |  |
| FHp23 | TCTTCTAACAACAGGCAACAGACAAGTCACACAAAACAAAAGCTATGTAAGGAAAACTGC | For construction of deletion template for clean deletion of DGA1 in YL W29 strain |
| FHp24 | TCAAAAAAATACTCATTAGCTATTTGCCTAACCCAGGCAGTTTTCCTTACATAGCTTTTG |  |
| FHp25 | TTCTCACGACCAATTACGTAC | For colony PCR of deletion of DGA1 in YL W29 strain |
| FHp26 | CAGTATTTATTCCGCTCACTC |  |
| FHp27 | TTCGATTCCGGGTCGGCGCAGAAAGTCCAGAAGAAACCTGGTTTTA | For cloning crRNA-DGA2_W29 to NATex_CrisprCas9-yl |
| FHp28 | GCTCTAAAACCAGGTTTCTTCTGGACTTTCTGCGCCGACCCGGAAT |  |
| FHp29 | ACACCATAGCTACACCCAAAGTCGCTGTACCCGTACCGCCCGCAAAATGTAGCATAACAC | For construction of deletion template for clean deletion of DGA2 in YL W29 strain |
| FHp30 | AATGTCAATATTATTTATCACTGTAAAGGCTACTGATGAGTGTTATGCTACATTTTGCGG |  |
| FHp31 | AACTCGACTGGAACCAAGTTC | For colony PCR of deletion of DGA2 in YL W29 strain |
| FHp32 | ACGAACAGGTTCAACATATGCG |  |
| FHp33 | TTCGATTCCGGGTCGGCGCAACTCAACGCCAAGTACCCGGGTTTTA | For cloning crRNA-LRO1_W29 to HPHex_CrisprCas9-yl |
| FHp34 | GCTCTAAAACCCGGGTACTTGGCGTTGAGTTGCGCCGACCCGGAAT |  |
| FHp35 | GTTGATCACAACCATCAATGACACAACCTGTGAATCGGAAGGCGACTGTCTAGAGAGTC | For construction of deletion template for clean deletion of LRO1 in YL W29 strain |
| FHp36 | TCCATCAGTGTGATATACAAAACATATTACTCTACAAAACGGACTCTCTAGACAGTCGC |  |
| FHp37 | AAACAAGACCATTCTGACGCG | For colony PCR of deletion of LRO1 in YL W29 strain |
| FHp38 | GACAGGAGTGGTTGTATATTGG |  |
| FHp39 | TTCGATTCCGGGTCGGCGCATGATAGGACCAGAGTCACCGGTTTTA | For cloning crRNA-FAO1_W29 to NATex_CrisprCas9-yl, to be used with clean deletion template |
| FHp40 | GCTCTAAAACCGGTGACTCTGGTCCTATCATGCGCCGACCCGGAAT |  |
| FHp41 | ATCAAAAAAAAGCTGTACACGCTTTGCATTAGCTTGATTACTTGACAATGTAAGCAAGAG | For construction of deletion template for clean deletion of FAO1 in YL W29 strain |
| FHp42 | AAATCATATACATTAGCATTACTTCTATTTCTAAACTCGTCTCTTGCTTACATTGTCAAG |  |
| FHp43 | TTCGATTCCGGGTCGGCGCATGGTGTTGACCCAGTTACCGGTTTTA | For cloning crRNA-FAO1_W29 to NATex_CrisprCas9-yl, to be used with dual cut system |
| FHp44 | GCTCTAAAACCGGTAACTGGGTCAACACCATGCGCCGACCCGGAAT |  |
| FHp45 | TTCGATTCCGGGTCGGCGCACTTCGAGGAAATGTCCATCGGTTTTA | For cloning crRNA-FAO1_W29 to HPHex_CrisprCas9-yl |
| FHp46 | GCTCTAAAACCGATGGACATTTCCTCGAAGTGCGCCGACCCGGAAT |  |
| FHp47 | AACAGCACAAAGTCGTTTCTCG | For colony PCR of deletion of FAO1 in YL W29 strain |
| FHp48 | ATCGAGATCGGCCTCTGATTAG |  |
| FHp49 | TTCGATTCCGGGTCGGCGCAAAAGGAACCCGAGGTCACAGGTTTTA | For cloning crRNA-CEX1_W29 to NATex_CrisprCas9-yl |
| FHp50 | GCTCTAAAACCTGTGACCTCGGGTTCCTTTTGCGCCGACCCGGAAT |  |
| FHp51 | TTCGATTCCGGGTCGGCGCACCTCACTCTCCTAACTCCAGGTTTTA | For cloning crRNA-CEX1_W29 to HPHex_CrisprCas9-yl |
| FHp52 | GCTCTAAAACCTGGAGTTAGGAGAGTGAGGTGCGCCGACCCGGAAT |  |
| FHp53 | ATTATCACCACTTGGGTGCATC | For colony PCR of deletion of CEX1 in YL W29 strain |
| FHp54 | GTTAGTGTTAAGGCAAGGAATTG |  |
| FHp55 | TTCGATTCCGGGTCGGCGCAAGGTCCAGACATGATTGAGGGTTTTA | For cloning crRNA-MDH2_W29 to NATex_CrisprCas9-yl |
| FHp56 | GCTCTAAAACCCTCAATCATGTCTGGACCTTGCGCCGACCCGGAAT |  |
| FHp57 | TTCGATTCCGGGTCGGCGCAGGAGACCCCTCCGAGCTCTGGTTTTA | For cloning crRNA-MDH1_W29 to HPHex_CrisprCas9-yl |
| FHp58 | GCTCTAAAACCAGAGCTCGGAGGGGTCTCCTGCGCCGACCCGGAAT |  |
| FHp59 | CCAAGATACCACACCATCATGG | For colony PCR of deletion of MDH2 in YL W29 strain |
| FHp60 | TTGACATTCATTGGCAATTCGG |  |
| FHp61 | TTCGATTCCGGGTCGGCGCAAGTCTCACCGAAACCAACCAGTTTTA | For cloning crRNA-MFE1_W29 to NATex_CrisprCas9-yl |
| FHp62 | GCTCTAAAACTGGTTGGTTTCGGTGAGACTTGCGCCGACCCGGAAT |  |
| FHp63 | TTCGATTCCGGGTCGGCGCAAGTACCTTGAAATCCGACAGGTTTTA | For cloning crRNA-MFE1_W29 to HPHex_CrisprCas9-yl |
| FHp64 | GCTCTAAAACCTGTCGGATTTCAAGGTACTTGCGCCGACCCGGAAT |  |
| FHp65 | CGGGACAACGAAACCATAAACC | For colony PCR of deletion of MFE1 in YL W29 strain |
| FHp66 | GTACTGTACTTGCAGTACACAG |  |
| FHp67 | TTCGATTCCGGGTCGGCGCAGTACAAGGAGGAGCTGGAGAGTTTTA | For cloning crRNA-PEX10_W29 to NATex_CrisprCas9-yl |
| FHp68 | GCTCTAAAACTCTCCAGCTCCTCCTTGTACTGCGCCGACCCGGAAT |  |
| FHp69 | TTCGATTCCGGGTCGGCGCAGCGCCGGCATGTACGCCATGGTTTTA | For cloning crRNA-PEX10_W29 to HPHex_CrisprCas9-yl |
| FHp70 | GCTCTAAAACCATGGCGTACATGCCGGCGCTGCGCCGACCCGGAAT |  |
| FHp71 | GTTGTCGATTGAATTGGTCGTG | For colony PCR of deletion of PEX10 in YL W29 strain |
| FHp72 | AAAATGCCAACTGAAATTTG |  |
| FHp73 | TTCGATTCCGGGTCGGCGCATGAAAGCAGAGATGGACGAGGTTTTA | For cloning crRNA-ARE1_W29 to NATex_CrisprCas9-yl |
| FHp74 | GCTCTAAAACCTCGTCCATCTCTGCTTTCATGCGCCGACCCGGAAT |  |
| FHp75 | TTCGATTCCGGGTCGGCGCAGGCAAACACCATCCACAGCAGTTTTA | For cloning crRNA-ARE1_W29 to HPHex_CrisprCas9-yl |
| FHp76 | GCTCTAAAACTGCTGTGGATGGTGTTTGCCTGCGCCGACCCGGAAT |  |
| FHp77 | TGACACGGCTCAGATTTGGTT | For colony PCR of deletion of ARE1 in YL W29 strain |
| FHp78 | GTGGCTCTACAGTGTCTCGTT |  |

**Table S2.** List of strains used and developed in this study.

| Strain | Description | Reference |
| --- | --- | --- |
| W29 | ATCC 20460 |  |
| FAOH1 | W29::A08-4UAS_PTEF-mhFAR-LIP2 | This study |
| FAOH2 | W29::A08-4UAS_PTEF-mhFAR-LIP2*dga1*Δ | This study |
| FAOH3 | W29::A08-4UAS_PTEF-mhFAR-LIP2*dga1*Δ*dga2*Δ | This study |
| FAOH4 | W29::A08-4UAS_PTEF-mhFAR-LIP2*lro1*Δ | This study |
| FAOH5 | W29::A08-4UAS_PTEF-mhFAR-LIP2*dga1*Δ*dga2*Δ*lro1*Δ | This study |
| FAOH6 | W29::A08-4UAS_PTEF-mhFAR-LIP2*fao1*Δ | This study |
| FAOH7 | W29::A08-4UAS_PTEF-mhFAR-LIP2*dga1*Δ*dga2*Δ*lro1*Δ*fao1*Δ | This study |
| FAOH8 | W29::A08-4UAS_PTEF-mhFAR-LIP2*dga1*Δ*dga2*Δ*lro1*Δ*fao1*Δ*cex1*Δ | This study |
| FAOHS1 | W29::A08-4UAS_PTEF-mhFAR-LIP2*dga1*Δ*dga2*Δ*lro1*Δ*are1*Δ | This study |
| FAOHS2 | W29::A08-4UAS_PTEF-mhFAR-LIP2*dga1*Δ*dga2*Δ*lro1*Δ*fao1*Δ*pex10*Δ | This study |
| FAOHS3 | W29::A08-4UAS_PTEF-mhFAR-LIP2*dga1*Δ*dga2*Δ*lro1*Δ*fao1*Δ*mfe1*Δ | This study |
| FAOHS4 | W29::A08-4UAS_PTEF-mhFAR-LIP2*dga1*Δ*dga2*Δ*lro1*Δ*fao1*Δ*mdh2*Δ | This study |
| FAOHS5 | W29::A08-4UAS_PTEF-mhFAR-LIP2*dga1*Δ*dga2*Δ*lro1*Δ*fao1*Δ*cex1*Δ*mdh2*Δ | This study |

**Table S3:** List of plasmids used in this study.

| Plasmid | Description | Source |
| --- | --- | --- |
| ***pSB1A3 (-BsaI)::rfp cassette*** | *Destination vector for golden gate assemblies* | **Addgene Plasmid #120730** |
| ***pCR4Blunt-TOPO- P2 TEF-4AUS*** | Topo vector harboring constitutive promoter for mhFAR expression | **Addgene Plasmid #120751** |
| ***pCR4Blunt-TOPO-(C)PTEF-4UAS(D2)*** | pCR4Blunt-TOPO- P2 TEF-4AUS cloned with appropriate overhangs | **This study** |
| ***pCR4Blunt-TOPO-T3 Lip2*** | Topo vector harboring terminator for mhFAR expression | **Addgene Plasmid #120774** |
| ***pCR4Blunt-TOPO-(E)TLip2(F)*** | pCR4Blunt-TOPO-T3 Lip2 cloned with appropriate overhangs | **This study** |
| ***pCR4Blunt-TOPO-(B)hph(C)*** | Topo vector harboring hygromycin resistance selection marker | **Addgene Plasmid #120736** |
| ***pCR4Blunt-TOPO-(A)A08_W29_LHB-NotI_Up(B)*** | Topo vector harboring A08 LHB for integration to A08 locus of YL W29 strain using crispr system, BsaI sites removed with site-directed mutagenesis | **This study** |
| ***pCR4Blunt-TOPO-(F)A08_W29_RHB-NotI_Down(M)*** | Topo vector harboring A08 RHB for integration to A08 locus of YL W29 strain using crispr system | **This study** |
| ***pCR4Blunt-TOPO-(D2)mhFAR(E)*** | Topo vector harboring YL codon optimized mhFAR gene | **This study** |
| ***NATex_CrisprCas9-yl_RFP*** | CRISPR-Cas9 YL destination plasmid, containing nourseothricin resistance selection marker | **Addgene Plasmid #129660** |
| ***HPHex_CrisprCas9-yl_RFP*** | CRISPR-Cas9 YL destination plasmid, containing hygromycin resistance selection marker | **Addgene Plasmid #129659** |
| ***pMCS-UAS1B16-TEF-Cre*** | Cre recombinase expressing plasmid to cure markers flanked by loxRP sites in YL, containing nourseothricin resistance selection marker | **Blazeck et al., 2013** |
| ***pSB1A3::A08-NotI_UP-hph-4UAS_PTEF-mhFAR-LIP2-A08-NotI_Down*** | Golden gate destination vector harboring YL expression casette for mhFAR, linearized at NotI cut sites before integration | **This study** |
| ***NATex_CrisprCas9-yl-sgRNA-A08_W29*** | NATex_CrisprCas9-yl containing sgRNA for A08 integration locus at YL W29 genome | **This study** |
| ***NATex_CrisprCas9-yl-sgRNA-DGA1_W29*** | NATex_CrisprCas9-yl containing sgRNA for DGA1 gene at YL W29 genome, used with clean deletion template | **This study** |
| ***NATex_CrisprCas9-yl-sgRNA-DGA2_W29*** | NATex_CrisprCas9-yl containing sgRNA for DGA2 gene at YL W29 genome, used with clean deletion template | **This study** |
| ***HPHex_CrisprCas9-yl-sgRNA-LRO1_W29*** | HPHex_CrisprCas9-yl containing sgRNA for LRO1 gene at YL W29 genome, used with clean deletion template | **This study** |
| ***NATex_CrisprCas9-yl-sgRNA-FAO1_W29*** | NATex_CrisprCas9-yl containing sgRNA for FAO1 gene at YL W29 genome, used with clean deletion template, used for FAOH8 strain | **This study** |
| ***NATex_CrisprCas9-yl-sgRNA-FAO1_W29*** | NATex_CrisprCas9-yl containing sgRNA for FAO1 gene at YL W29 genome, targets left flank, used for FAOH9 strain | **This study** |
| ***HPHex_CrisprCas9-yl-sgRNA-FAO1_W29*** | HPHex_CrisprCas9-yl containing sgRNA for FAO1 gene at YL W29 genome, targets right flank, used for FAOH9 strain | **This study** |
| ***NATex_CrisprCas9-yl-sgRNA-CEX1_W29*** | NATex_CrisprCas9-yl containing sgRNA for CEX1 gene at YL W29 genome, targets left flank | **This study** |
| ***HPHex_CrisprCas9-yl-sgRNA-CEX1_W29*** | HPHex_CrisprCas9-yl containing sgRNA for CEX1 gene at YL W29 genome, targets right flank | **This study** |
| ***NATex_CrisprCas9-yl-sgRNA-MDH2_W29*** | NATex_CrisprCas9-yl containing sgRNA for MDH2 gene at YL W29 genome, targets left flank | **This study** |
| ***HPHex_CrisprCas9-yl-sgRNA-MDH2_W29*** | HPHex_CrisprCas9-yl containing sgRNA for MDH2 gene at YL W29 genome, targets right flank | **This study** |
| ***NATex_CrisprCas9-yl-sgRNA-MFE1_W29*** | NATex_CrisprCas9-yl containing sgRNA for MFE1 gene at YL W29 genome, targets left flank | **This study** |
| ***HPHex_CrisprCas9-yl-sgRNA-MFE1_W29*** | HPHex_CrisprCas9-yl containing sgRNA for MFE1 gene at YL W29 genome, targets right flank | **This study** |
| ***NATex_CrisprCas9-yl-sgRNA-PEX10_W29*** | NATex_CrisprCas9-yl containing sgRNA for PEX10 gene at YL W29 genome, targets left flank | **This study** |
| ***HPHex_CrisprCas9-yl-sgRNA-PEX10_W29*** | HPHex_CrisprCas9-yl containing sgRNA for PEX10 gene at YL W29 genome, targets right flank | **This study** |
| ***NATex_CrisprCas9-yl-sgRNA-ARE1_W29*** | NATex_CrisprCas9-yl containing sgRNA for ARE1 gene at YL W29 genome, targets right flank | **This study** |
| ***HPHex_CrisprCas9-yl-sgRNA-ARE1_W29*** | HPHex_CrisprCas9-yl containing sgRNA for ARE1 gene at YL W29 genome, targets left flank | **This study** |

**Table S4.** *mhFAR* (*Marinobacter hydrocarboniclasticus*) nucleotide sequence, codon optimized for *Yarrowia lipolytica.*

| Gene name | Sequence |
| --- | --- |
| ***mhFAR*** | ATGGCCATCCAGCAGGTGCATCACGCCGATACATCTTCTAGCAAGGTCCTCGGCCAGCTCAGAGGAAAGCGAGTTCTCATCACAGGCACAACCGGATTCCTGGGTAAGGTCGTTCTGGAACGACTCATTCGTGCCGTGCCTGATATTGGTGCAATTTACCTGCTCATCAGAGGTAACAAGCGGCATCCCGACGCTCGGTCCCGATTTCTGGAGGAAATTGCTACCAGCTCTGTCTTCGATCGACTGAGAGAGGCTGACTCCGAAGGATTTGACGCCTTTCTTGAGGAGCGAATCCACTGTGTCACTGGTGAGGTCACCGAGGCTGGCTTTGGAATTGGCCAGGAGGATTACCGGAAGCTTGCTACTGAACTCGATGCTGTGATTAACTCTGCCGCCTCGGTGAACTTCCGGGAGGAACTTGACAAGGCTCTGGCCATTAACACGCTCTGCCTTCGTAATATCGCTGGAATGGTCGACCTGAACCCAAAACTCGCCGTCCTCCAGGTTTCTACCTGCTACGTTAACGGCATGAACAGCGGTCAAGTGACAGAGTCTGTTATCAAGCCCGCTGGAGAAGCAGTCCCTCGATCGCCCGACGGCTTCTACGAGATCGAGGAGCTTGTCCGACTCCTGCAGGACAAGATCGAGGATGTCCAGGCCCGATACTCCGGCAAGGTGCTGGAGCGAAAGCTCGTCGATCTCGGAATCCGAGAAGCTAACCGTTACGGATGGAGCGATACCTACACCTTCACCAAGTGGCTCGGCGAACAGCTTCTGATGAAGGCCCTCAACGGCCGAACCCTTACAATTCTTCGACCCTCTATTATCGAGTCTGCTCTCGAGGAGCCCGCTCCTGGCTGGATCGAGGGAGTGAAGGTTGCTGATGCCATCATCCTCGCCTACGCTCGAGAGAAGGTGACTCTGTTCCCTGGAAAGCGATCTGGCATTATTGACGTCATCCCTGTCGACCTGGTCGCCAACTCTATTATTCTTAGCCTTGCCGAGGCCCTGGGCGAACCCGGACGACGTCGAATTTACCAGTGCTGCTCCGGTGGCGGAAACCCTATCAGCCTGGGTGAGTTCATCGATCATCTCATGGCCGAGAGCAAGGCCAACTACGCCGCTTACGACCACCTGTTTTATCGACAGCCTTCCAAGCCCTTCCTTGCTGTTAACCGAGCCCTGTTCGATCTGGTGATCTCTGGCGTTAGACTCCCTCTCTCTCTCACCGATCGAGTGCTCAAGCTCCTCGGCAACTCCCGAGAtCTTAAGATGCTTCGTAACCTTGATACAACCCAATCCCTGGCCACCATCTTTGGATTCTACACCGCTCCCGACTACATCTTCCGAAACGACGAGCTCATGGCTCTCGCTAACCGGATGGGAGAGGTCGACAAAGGTCTGTTCCCCGTGGATGCCCGACTTATCGACTGGGAGCTCTACCTGCGAAAGATCCATCTGGCTGGCCTGAACCGATACGCACTGAAGGAGCGTAAGGTTTACTCCCTGAAGACTGCCCGACAGCGAAAGAAGGCCGCTTAA |

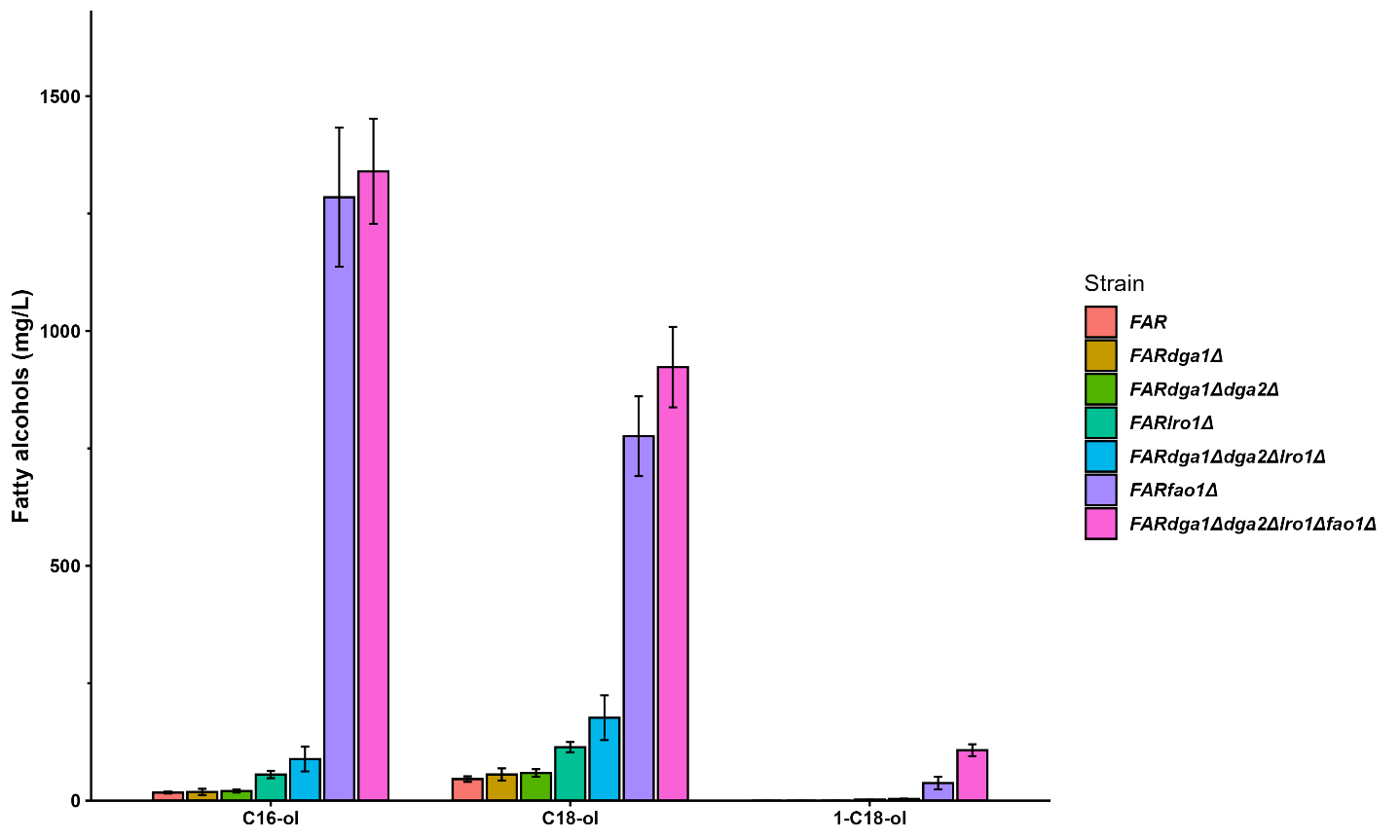

**Figure S1.** **Fatty alcohol chain-length profile at peak titers for studied strains.** Growth experiments were performed in shake flasks containing YNB medium with 40 g/L glucose (C/N 5) using shake flasks. Captured at 72h. Error bars denote the standard deviation from three biological replicates

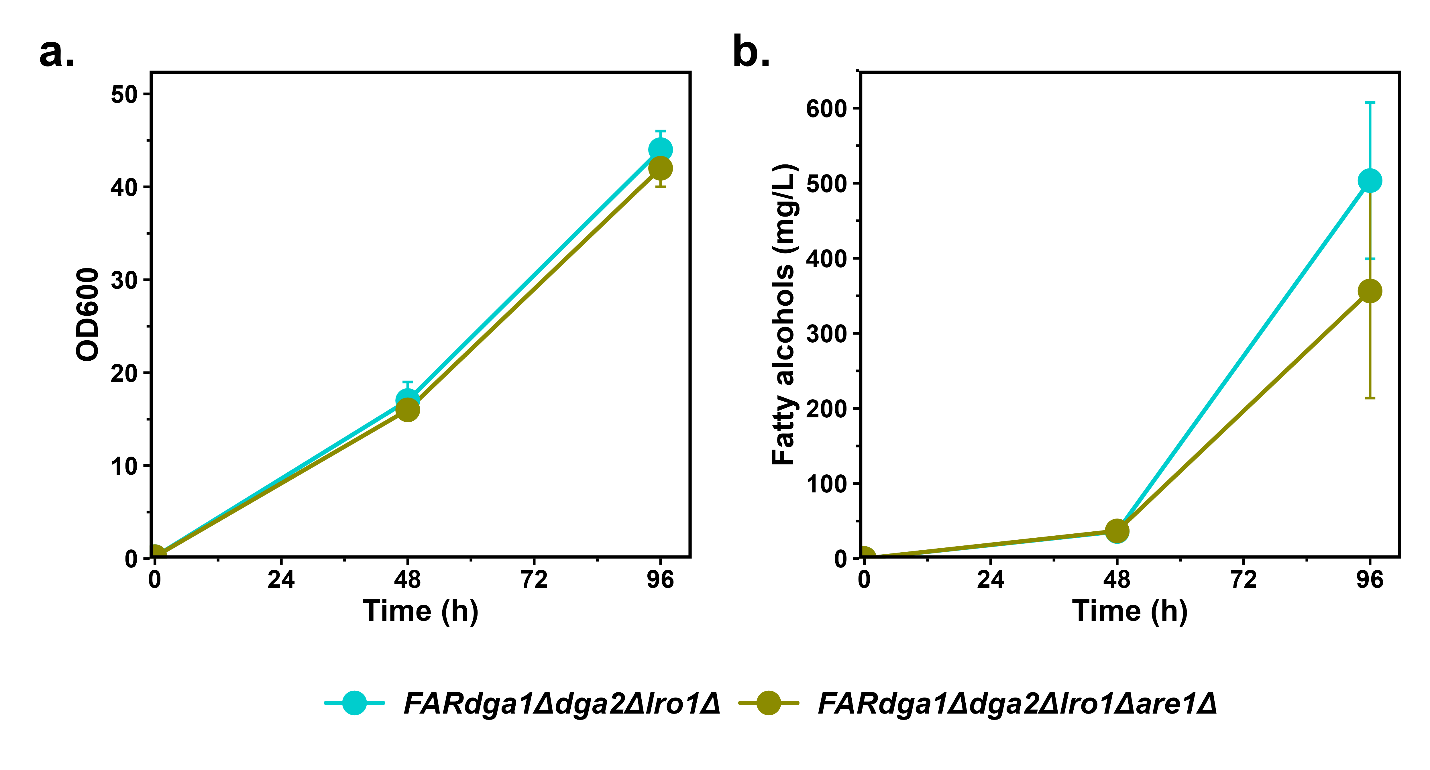

**Figure S2.** **Effect of *ARE1* deletion.** Growth experiments were performed in shake flasks containing YNB medium with 40 g/L glucose (C/N 5) using shake flasks. Legend: (a.) cell growth (OD600), (b.) total fatty alcohol production. Error bars denote the standard deviation from two biological replicates.

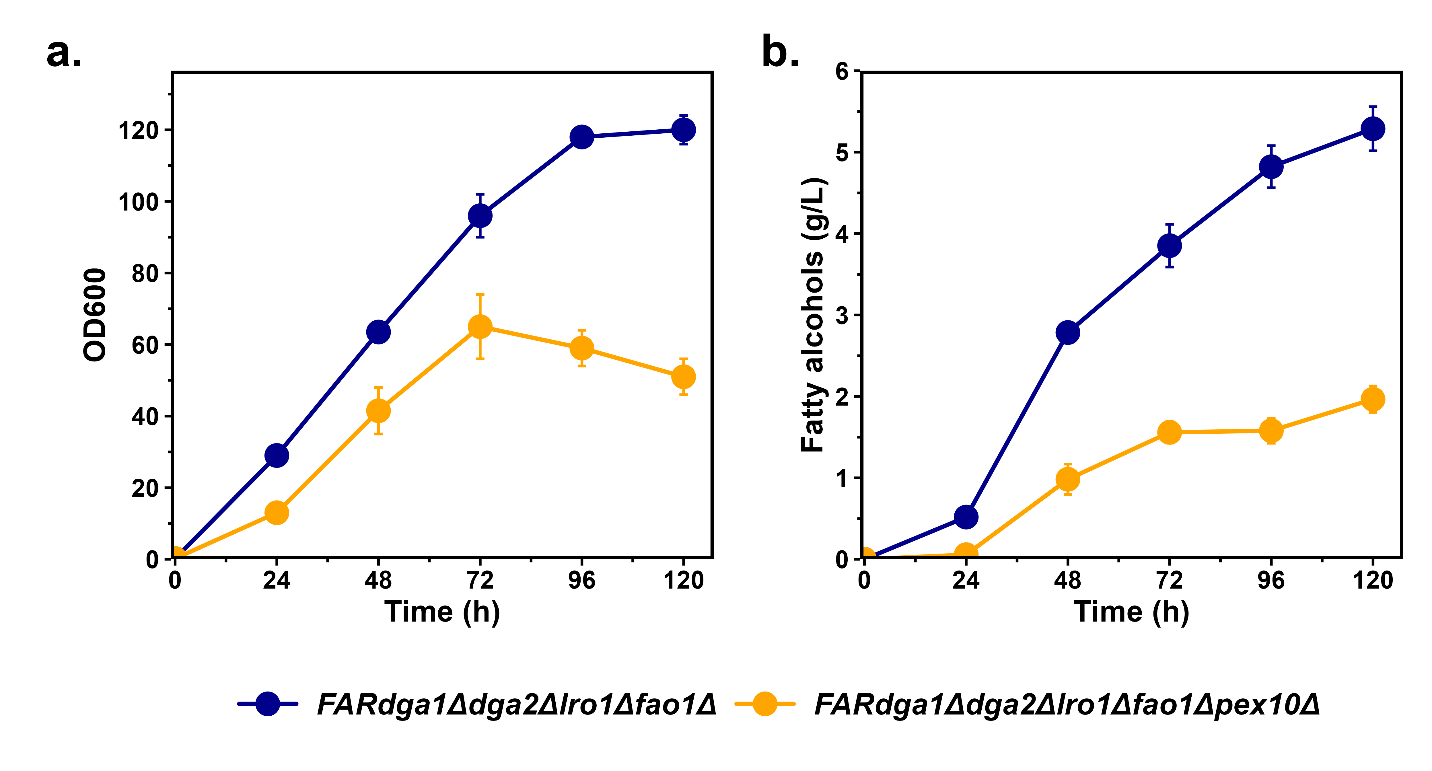

**Figure S3.** **Effect of *PEX10* deletion.** Growth experiments were performed in shake flasks containing YNB medium with 80 g/L glucose (C/N 10), supplemented with 1.92 g/L yeast synthetic drop-out medium supplements without tryptophan to support *PEX10* deficient culture growth, using shake flasks. Legend: (a.) cell growth (OD600), (b.) total fatty alcohol production. Error bars denote the standard deviation from two biological replicates.

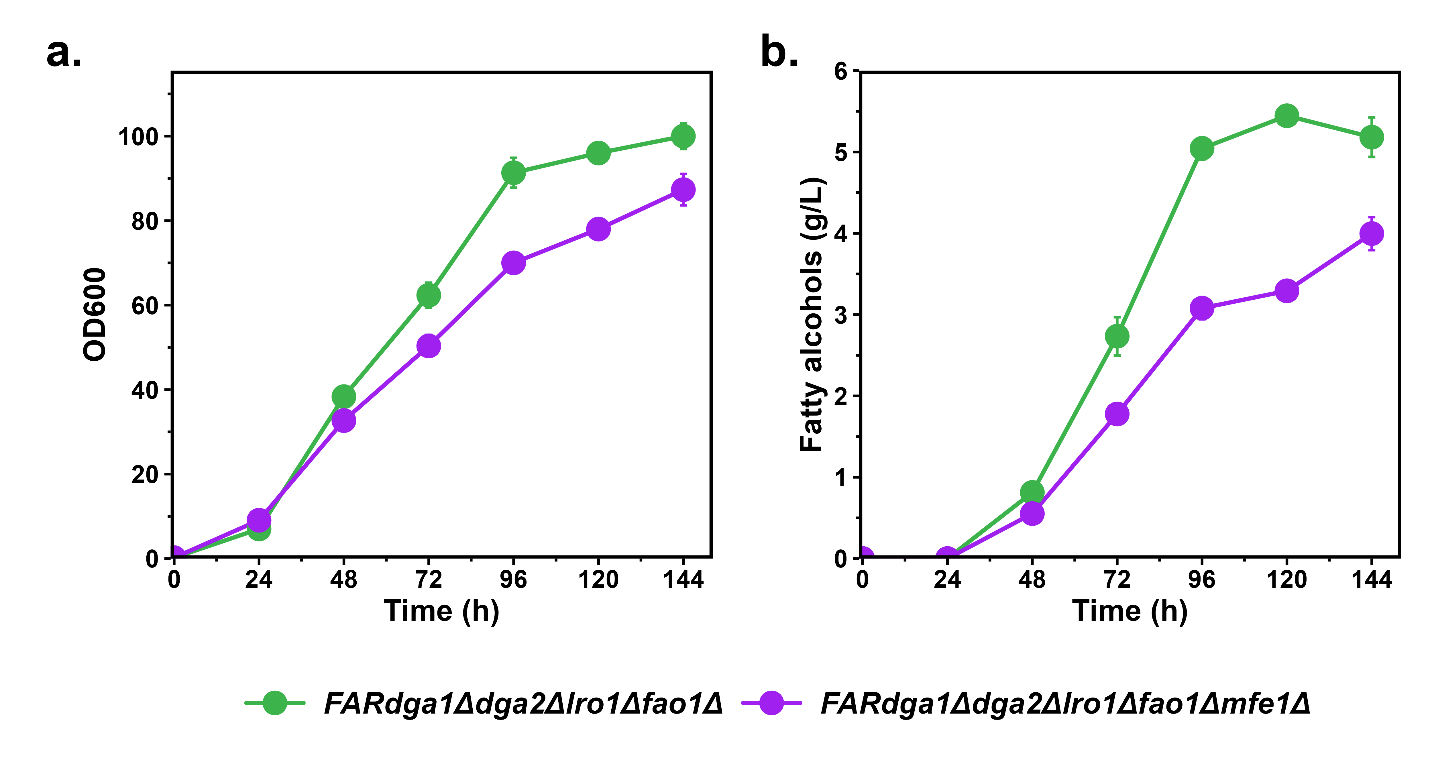
**Figure S4.** **Effect of *MFE1* deletion.** Growth experiments were performed in shake flasks containing YNB medium with 80 g/L glucose (C/N 10) using shake flasks. Legend: (a.) cell growth (OD600), (b.) total fatty alcohol production. Error bars denote the standard deviation from two biological replicates.

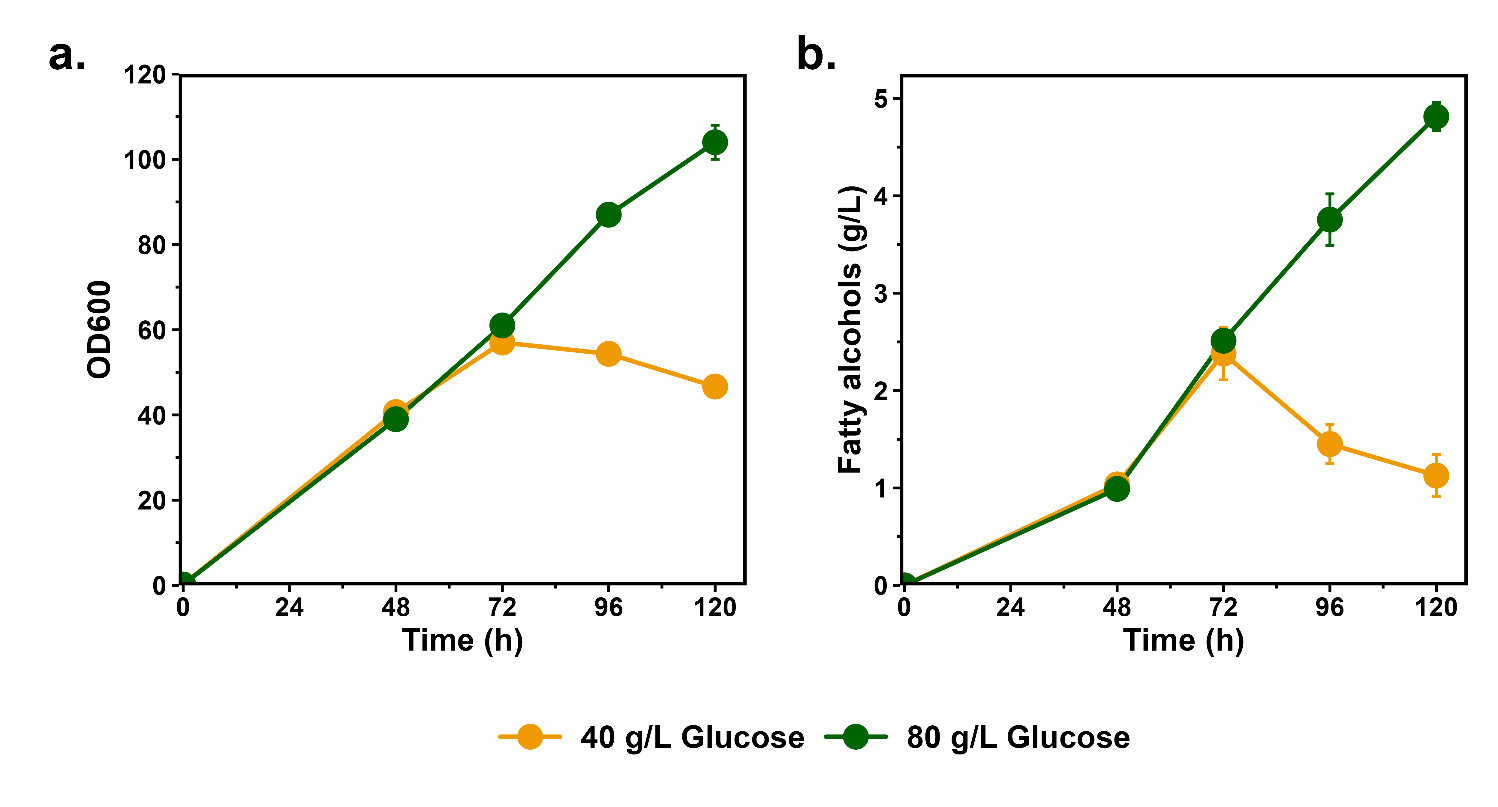

**Figure S5.** **Effect of doubled glucose concentration on fatty alcohol production.** Growth experiments were performed using the *mhFAR*-expressing *dga1*Δ*dga2*Δ*lro1*Δ*fao1*Δ strain in shake flasks containing YNB medium with 40 g/L glucose and a C/N ratio of 5 or 80 g/L glucose and a C/N ratio of 10, respectively. Legend: (a.) cell growth (OD600), (b.) total fatty alcohol production. Error bars denote the standard deviation from two biological replicates.

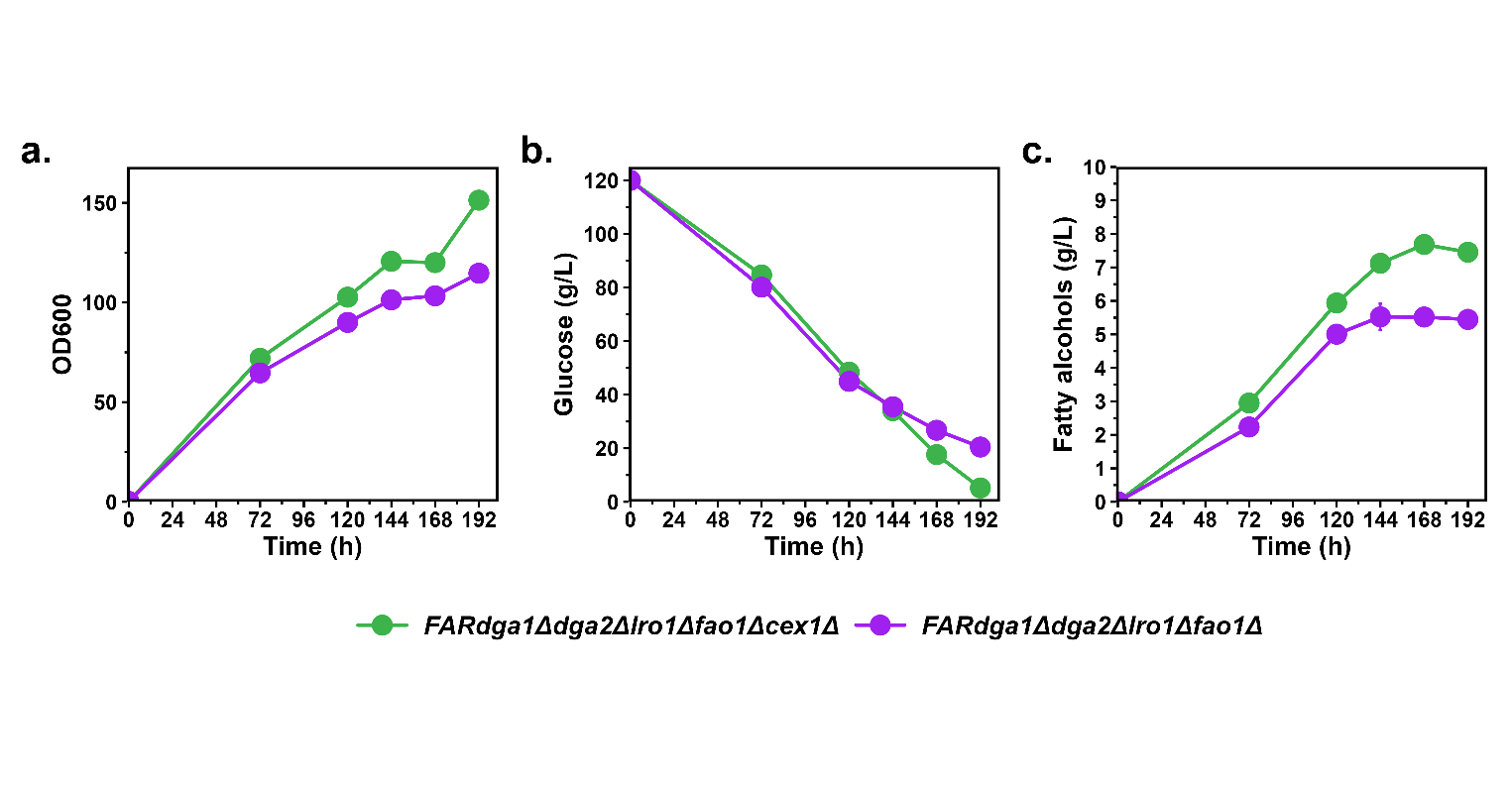
**Figure S6.** **Effect of tripled glucose concentration on fatty alcohol production.** Growth experiments were performed in shake flasks containing YNB medium with 120 g/L glucose (C/N 60) using shake flasks. Legend: (a.) cell growth (OD600), (b.) glucose consumption, (c.) total fatty alcohol production. Error bars denote the standard deviation from three biological replicates.

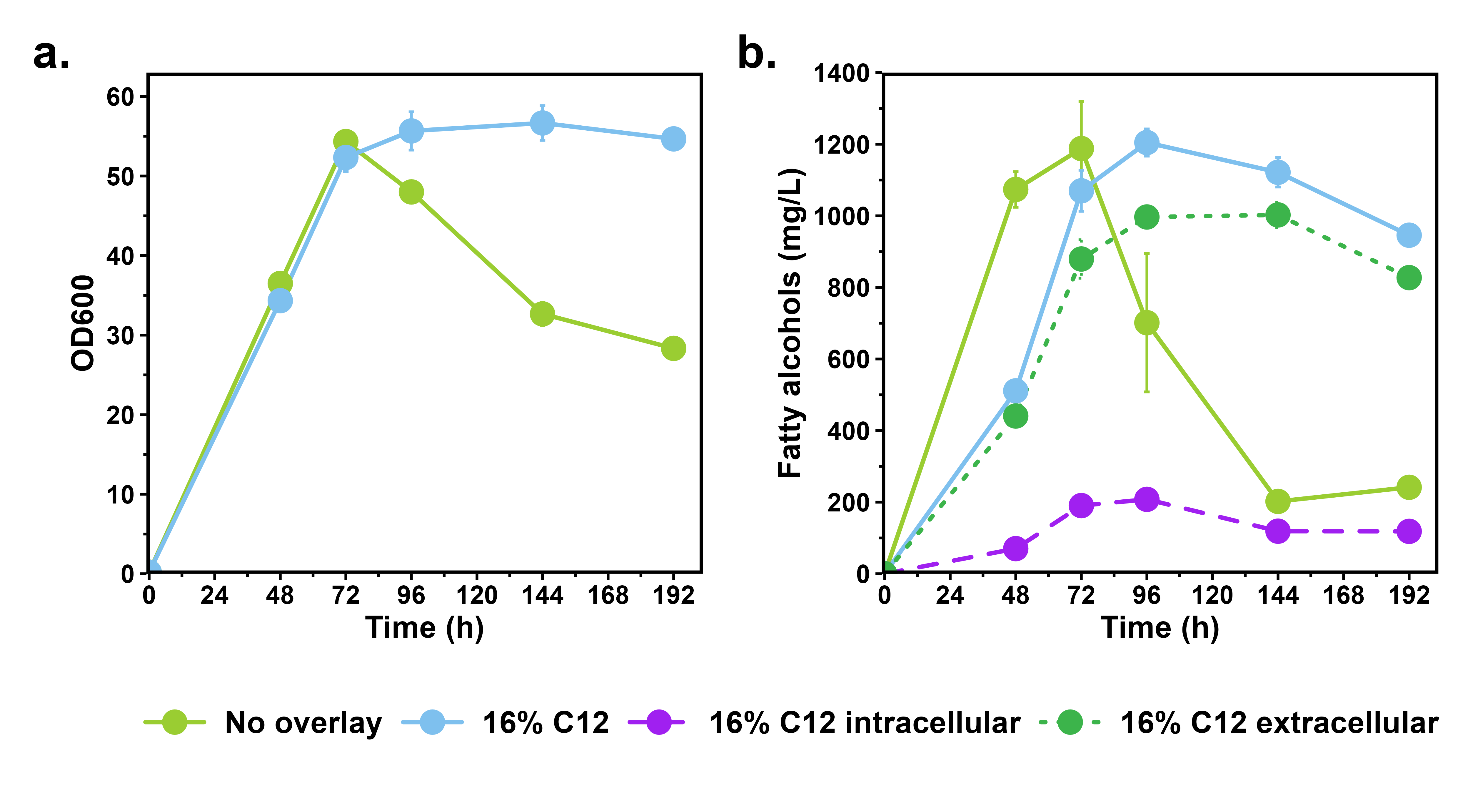

**Figure S7.** **Effect of dodecane on fatty alcohol production.** Growth experiments were performed using the *mhFAR*-expressing *dga1Δdga2Δlro1Δfao1Δ* strain in shake flasks containing YNB medium with 40 g/L glucose and a C/N ratio of 5. Legend: (a.) cell growth (OD600) and (b.) total fatty alcohol production. Error bars denote the standard deviation from two biological replicates. Intracellular and extracellular portions of overlay cultures are shown in dashed and dotted lines, respectively.

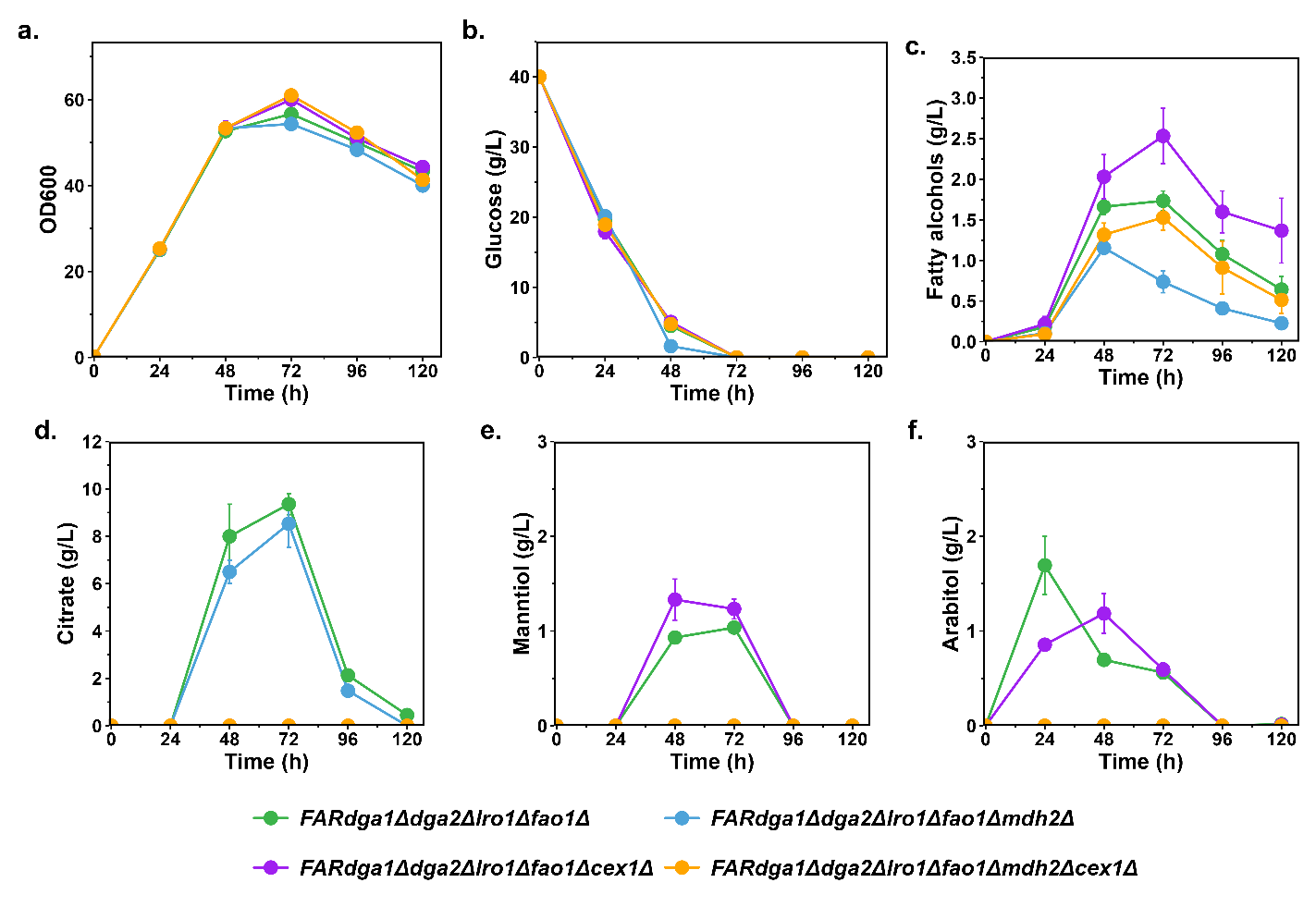

**Figure S8.** **Effect of *MDH2* deletion.** Growth experiments were performed in shake flasks containing YNB medium with 40 g/L glucose (C/N 5) using shake flasks. Legend: (a.) cell growth (OD600), (b.) glucose consumption, (c.) total fatty alcohol production, (d.) citrate production, (e.) mannitol production, and (f.) arabitol production. Error bars denote the standard deviation from three biological replicates.

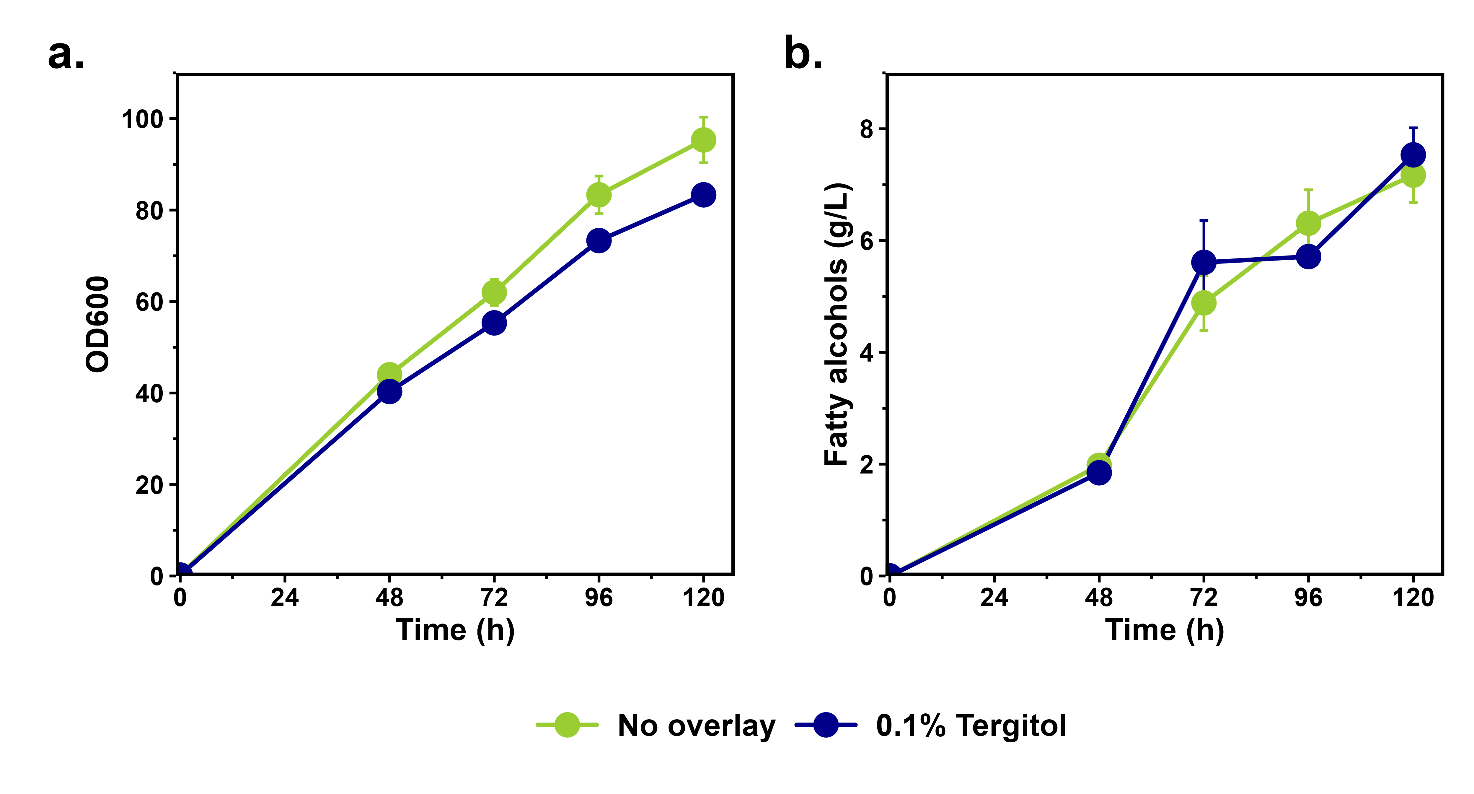

**Figure S9.** **Effect of 0.1% tergitol on fatty alcohol production.** Growth experiments were performed using the *mhFAR*-expressing *dga1*Δ*dga2*Δ*lro1*Δ*fao1*Δ*cex1*Δ strain in shake flasks containing YNB medium with 80 g/L glucose and a C/N ratio of 60. Legend: (a.) cell growth (OD600) and (b.) total fatty alcohol production. Error bars denote the standard deviation from three biological replicates
